# Nuclear Factor-Y is a Pervasive Regulator of Neuronal Gene Expression

**DOI:** 10.1101/2023.02.14.528575

**Authors:** Pedro Moreira, Paul Papatheodorou, Shuer Deng, Sandeep Gopal, Ava Handley, David Powell, Roger Pocock

## Abstract

Nervous system function relies on the establishment of complex gene expression programs that provide neuron-type-specific and core pan-neuronal features. These complementary regulatory paradigms are controlled by terminal selector and parallel-acting transcription factors (TFs), respectively. Here, we identify the Nuclear Factor Y (NF-Y) TF as a pervasive regulator of both neuron-type-specific and pan-neuronal gene expression. Mapping global NF-Y targets reveals direct binding to the *cis*-regulatory regions of pan-neuronal genes and terminal selector TFs. We show that NFYA-1 controls pan-neuronal gene expression directly through binding to CCAAT boxes in target gene promoters and indirectly by regulating the expression of terminal selector TFs. Further, we find that NFYA-1 regulation of neuronal gene expression is important for neuronal activity and motor function. Thus, our research sheds light on how global neuronal gene expression programs are buffered through direct and indirect regulatory mechanisms.

## INTRODUCTION

A functional nervous system requires the correct expression of neuronal gene batteries that control both neuron-type-specific and pan-neuronal functions (Stefanakis et al., 2015). Neuron-type-specific genes encode neurotransmitter pathway components, ion channels, neuropeptides and receptors that provide specific sensory modalities and specialized neuronal features (Hobert, 2016). In contrast, genes expressed pan-neuronally encode universal neuronal functions, such as synaptic vesicle release, neuropeptide processing and neuronal housekeeping features (Taylor et al., 2021). Extensive genetic and *cis*-regulatory analyses in the *Caenorhabditis elegans* model have revealed distinct modes of regulation for neuron-type-specific and pan-neuronal genes. In general, neuron-type-specific gene expression is regulated by the non-redundant activity of terminal selector transcription factors (TFs) that bind discrete *cis-*regulatory elements (Glenwinkel et al., 2021; Hobert, 2011; Stefanakis et al., 2015). In contrast, pan-neuronal gene expression is controlled by redundantly acting TFs that bind diverse *cis-*regulatory elements (Stefanakis et al., 2015). The complementary regulatory paradigms controlling neuron-type-specific and pan-neuronal gene batteries may provide evolvability and robustness to gene expression, respectively. A recent study revealed that pan-neuronal gene expression is redundantly controlled by six members of the CUT homeodomain TF family in *C. elegans* (Leyva-Diaz and Hobert, 2022). These CUT TFs also cooperate with specific terminal selector TFs to control pan-neuronal expression in certain neurons, but do not control neuron-type-specific features (Leyva-Diaz and Hobert, 2022). It is unknown whether any TFs in the *C. elegans* nervous system can control both neuron-type-specific and pan-neuronal gene expression.

The NF-Y transcription factor (NF-Y), also known as the CCAAT-binding factor (CBF), is a highly conserved trimeric protein complex composed of NF-YA, NF-YB and NF-YC subunits (Nardini et al., 2013). NF-Y regulates gene expression by directly binding to CCAAT boxes that occur in approximately 30% of eukaryotic promoters (Dolfini et al., 2009). These CCAAT boxes are often located <100 nucleotides from transcriptional start sites where NF-Y recruits RNA polymerase II and basal transcription factors to regulate target gene expression (Kabe et al., 2005). The NF-YB and NF-YC subunits dimerize through their histone-fold domains (HFDs) prior to associating with NF-YA, which harbors DNA-binding and transactivation domains (Dolfini et al., 2009). Structural studies have shown that NF-YA makes sequence-specific DNA contacts, whereas NF-YB/NF-YC interact non-specifically with DNA through their HFDs (Nardini et al., 2013). NF-Y interacts with the minor groove of DNA to induce bending and chromatin accessibility (Nardini et al., 2013). NF-Y activity is essential for early mouse development as NF-YA knockout causes embryonic lethality (Bhattacharya et al., 2003). Studies have also shown that NF-Y controls cell proliferation of embryonic stem cells (ESCs) and hematopoietic stem cells (Bungartz et al., 2012; Dolfini et al., 2012). In ESCs, NF-Y regulates housekeeping genes by binding to their proximal promoters, and regulates genes required for cell identity by binding to distal enhancers in collaboration with cell type-specific master TFs (Oldfield et al., 2014; Tiwari et al., 2011). In the murine nervous system, conditional depletion of NF-YA in postmitotic neurons causes degeneration, possibly through dysregulation of endoplasmic reticulum homeostasis (Yamanaka et al., 2014). In *Drosophila*, a genetic screen also revealed a role for NF-YC in promoting synaptic connectivity of photoreceptor axons by repressing the key zinc-finger TF, Senseless (Morey et al., 2008). However, the precise mechanisms by which NF-Y controls gene expression in the developing nervous system is not well understood.

In this study, we reveal that the NF-Y TF complex regulates both neuron-type-specific and pan-neuronal gene expression in *C. elegans*. In a genetic screen for regulators of PVQ interneuron fate, we found that NF-Y is required for neuron-type-specific terminal identity and pan-neuronal features in this neuron. By mapping NFYA-1 ChIP-Seq data, we show that NFYA-1 directly binds to the *cis*-regulatory control regions of pan-neuronal factors, but has limited binding to neuron-type-specific gene promoters. By measuring the expression of transcriptional, translational, and endogenous reporters in specific *C. elegans* ventral nerve cord (VNC) motor neurons, we confirmed a critical role for NFYA-1 in controlling pan-neuronal gene expression. As one may predict from reduced pan-neuronal gene expression in motor neurons, loss of NFYA-1 causes deficits in neuronal activity and locomotion. We found that regulation of neuronal gene expression requires cell-autonomous expression of NFYA-1 and the presence of proximal CCAAT binding sites. Interestingly, despite the lack of NF-Y binding to neuron-type-specific genes, we detected changes in their expression in *nfya-1* mutant animals, suggesting an indirect mode of regulation. In support of this, we found that NFYA-1 also regulates the expression of specific terminal selector TFs that are required for correct neuron-type-specific gene expression. Our findings point to an additional layer of neuronal gene regulation governed by the NFYA-1 transcription factor, which, unlike previously identified factors, regulates both neuron-type-specific and pan-neuronal genes. Evidence of mouse NFY-A1 binding to orthologous pan-neuronal genes suggests that the NF-Y complex performs a conserved function in controlling neuronal gene expression across evolution.

## RESULTS

### The NF-Y/CCAAT Complex Regulates Interneuron Cell Fate

The glutamatergic PVQ interneurons extend axons along the entire *C. elegans* VNC to communicate temperature signals between the head and tail (Figure 1A) (Motomura et al., 2022). Mechanisms regulating PVQ axon guidance and outgrowth have been extensively studied (Norgaard et al., 2018; Pocock and Hobert, 2008; Schmitz et al., 2007); however, the control of PVQ fate specification is poorly defined. We performed a forward genetic screen using ethyl methanesulfonate (EMS) to isolate mutants with abnormal PVQ expression of the *sra-6prom::gfp* transgene, which is expressed in the PVQs and two pairs of head neurons (Figure 1A). We isolated the *rp120* mutant that exhibits reduced *sra-6prom::gfp* expression in the PVQs, without detectably affecting head neuron expression (Figure 1A). One-step whole-genome sequencing and single-nucleotide polymorphism mapping (Doitsidou et al., 2010) identified the *rp120* lesion as a mutation at the splice acceptor site of the fourth exon of the *nfyc-1* gene (Figure 1B). *nfyc-1* encodes a subunit of the trimeric NF-Y/CCAAT TF complex - composed of NF-YA, NF-YB, and NF-YC subunits (Figure 1C) (Nardini et al., 2013). We found that 95% of *nfyc-1(rp120)* mutant animals exhibited dim or undetectable expression of *sra-6prom::gfp* in the PVQs (Figure 1D). An independently derived *nfyc-1(tm4541)* deletion allele phenocopied the *nfyc-1(rp120)* mutation, demonstrating that *nfyc-1* loss causes defects in PVQ cell fate specification (Figure 1B, D).

**Figure 1.**
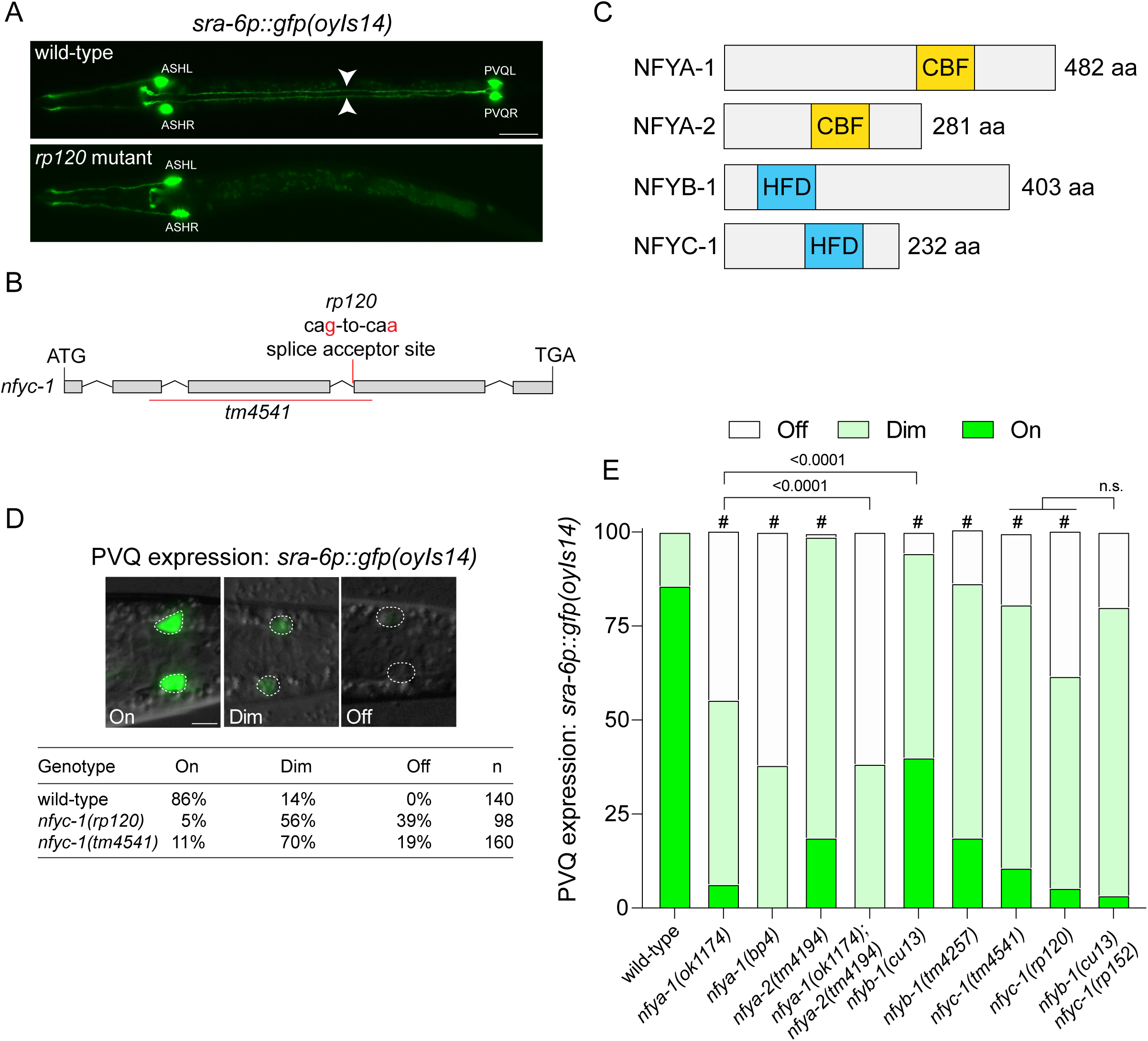
The NF-Y complex controls PVQ interneuron fate. (A) Fluorescent micrographs of *sra-6prom::gfp(oyIs14)* expression in wild-type and *rp120* mutant L1 larvae. In wild-type animals (top panel), the PVQL/R interneuron cell bodies (named PVQs from here) extend anteriorly-directed axons in the ventral nerve cord (white arrowheads) that terminate at the nerve ring. In *rp120* animals (bottom panel), *sra-6prom::gfp(oyIs14)* expression is undetectable in the PVQs. *sra-6prom::gfp(oyIs14)* expression in the ASH neurons is not affected by the *rp120* mutation (see Figure S2D for scoring). *sra-6prom::gfp(oyIs14)* is also weakly expressed in the ASI neurons and was not scored (out of field of view). Ventral views, anterior to the left. Scale bar: 10 μm. (B) *nfyc-1* genomic structure showing the *rp120* splice acceptor site mutation and *tm4541* deletion (red bar). (C) Protein domain organization of the NFY transcription factors in *C. elegans*. CBF = CCAAT-binding factor domain. HFD = histone fold domain. (D) Quantification of *sra-6prom::gfp(oyIs14)* expression in the PVQs of wild-type, *nfyc-1(rp120)* and *nfyc-1(tm4541)* animals. n refers to *sra-6prom::gfp(oyIs14)* expression per PVQ neuron. Scale bar: 10 μm. (E) *sra-6prom::gfp(oyIs14)* expression in the PVQs of wild-type, NF-Y complex single mutants and the following double mutants: *nfya-1(ok1174); nfya-2(tm4194)* and *nfyb-1(cu13); nfyc-1(rp152)*. Data presented as PVQ expression per neuron, as shown in (D). n = 60-160. Significance assessed by one-way ANOVA with Tukey’s correction. # = P<0.0001 compared to wild-type.

The *C. elegans* genome encodes two NFYA subunits (NFYA-1 and NFYA-2) and one each of NFYB-1 and NFYC-1 subunits. We surveyed the available NF-Y genetic mutants to assess their requirement to drive *sra-6p::gfp* expression (Figure 1E and S1). We found that loss-of-function mutations in all *nfy* genes resulted in decreased PVQ expression of *sra-6prom::gfp* (Figure 1E). Loss of *nfya-1* caused a significantly stronger reduction in *sra-6prom::gfp* expression compared to *nfya-2*, although the *nfya-1* mutant phenotype was enhanced by the loss of *nfya-2* (Figure 1E). This suggests that NFYA subunits can act in separate NF-Y complexes to control *sra-6prom::gfp* reporter expression. We found that *nfyb-1* and *nfyc-1* mutants also had reduced PVQ *sra-6prom::gfp* expression, although they had weaker phenotypes than the simultaneous loss of *nfya-1/2* (Figure 1E). Because *nfyb-1* and *nfyc-1* are closely linked on chromosome II, we used CRISPR-Cas9 editing to introduce a premature nonsense cassette into the first exon of *nfyc-1* in the *nfyb-1(cu13)* mutant background (Figure 1E and S1). This *nfyb-1(cu13) nfyc-1(rp152)* double mutant exhibited a similar decrease in *sra-6prom::gfp* expression in the PVQs as *nfyc-1* single mutants (Figure 1E). These data suggest that NFYA-1 and/or NFYA-2 may partially regulate *sra-6prom::gfp* expression independently of the NFYB/C subunits.

### NFYA-1 Cell-Autonomously Controls Interneuron Development

We examined the expression of NF-Y complex subunits using previously generated transcriptional reporters in which *nfya-1*, *nfyb-1* and *nfyc-1* promoters drive expression of mCherry fused to histone H1 (Figure S2A) (Sarov et al., 2006). These reporters are expressed broadly albeit with distinct domains of expression, likely due to the absence of complete regulatory information (Figure S2A). We crossed the *sra-6prom::gfp* transgene into these *nfy* reporter strains and found that all *nfy* promoters drive expression in the PVQ neurons (Figure S2A). This supports independent embryonic sequencing data that detected *nfy* gene expression in the PVQ neurons from birth (Packer et al., 2019). We focused on the DNA-binding NFYA-1 subunit to assess whether the NF-Y complex acts autonomously to control PVQ *sra-6prom::gfp* expression. We found that expressing a *nfya-1*-containing fosmid or *nfya-1* cDNA in the PVQs using the *npr-11* promoter restored *sra-6prom::gfp* PVQ expression to *nfya-1(ok1174)* animals (Figure S2B). As the PVQ neurons are born during embryogenesis, we investigated whether NFYA-1 is required to specify PVQ cell fate during the early stages of development. The *sra-6prom::gfp* transgene is only reliably detected from late embryonic stages, therefore, we compared the penetrance of PVQ *sra-6prom::gfp* expression in *nfya-1(ok1174)* animals at the L1 and L4 larval stages (Figure S2C) and found that the decrease in *sra-6prom::gfp* expression was identical at these two stages. Together, these data show that the NF-Y complex acts cell autonomously, likely during early development, to promote *sra-6prom::gfp* expression in the PVQ interneurons.

### NFYA-1 Regulates Neuron-Type-Specific and Pan-Neuronal Gene Expression in the PVQ Interneurons

To examine whether NFYA-1 plays a broader role in specifying PVQ neuron fate, we surveyed the expression of a battery of other PVQ identity genes (Figure 2). We crossed 8 PVQ-expressing reporter transgenes into *nfya-1(ok1174)* animals and found that 7 exhibited reduced expression in *nfya-1* mutants (Figure 2). We quantified the fluorescence intensity of these reporters in wild-type and *nfya-1(ok1174)* animals to fully determine the importance of the NF-Y complex in regulating PVQ cell fate (Figure 2A-H). We found that the expression of 4 reporters was almost completely abrogated (SRA-6, SRG-32, SRH-277, and SRI-1) and 3 others (DOP-1, EGL-47 and NPR-11) exhibited reduced expression – all of which encode transmembrane receptors (Figure 2A-G). In contrast, the expression of a reporter for the secreted immunoglobulin domain protein ZIG-5 is unaffected by *nfya-1* loss (Figure 2H). We found that all PVQ promoter transgenes that are affected by NFYA-1 loss contain multiple potential NF-Y binding sites (Figure S3A). To examine the functional relevance of NF-Y binding sites for PVQ expression, we focused on two genes (*sri-1* and *srh-277*) that are strongly regulated by NFYA-1 (Figure 2C-D). We mutated individual putative CCAAT binding sites in the *sri-1* and *srh-277* promoters and generated transgenic worms expressing these reporters (Figure S3B-C). We found that mutating individual NF-Y binding sites reduced GFP expression driven by the *sri-1* and *srh-277* promoters, suggesting direct regulation of gene expression by NFYA-1 (Figure S3B-C). These findings suggest that NFYA-1 directly regulates the expression of many neuron-type-specific features in PVQ neurons and likely in a direct manner.

**Figure 2.**
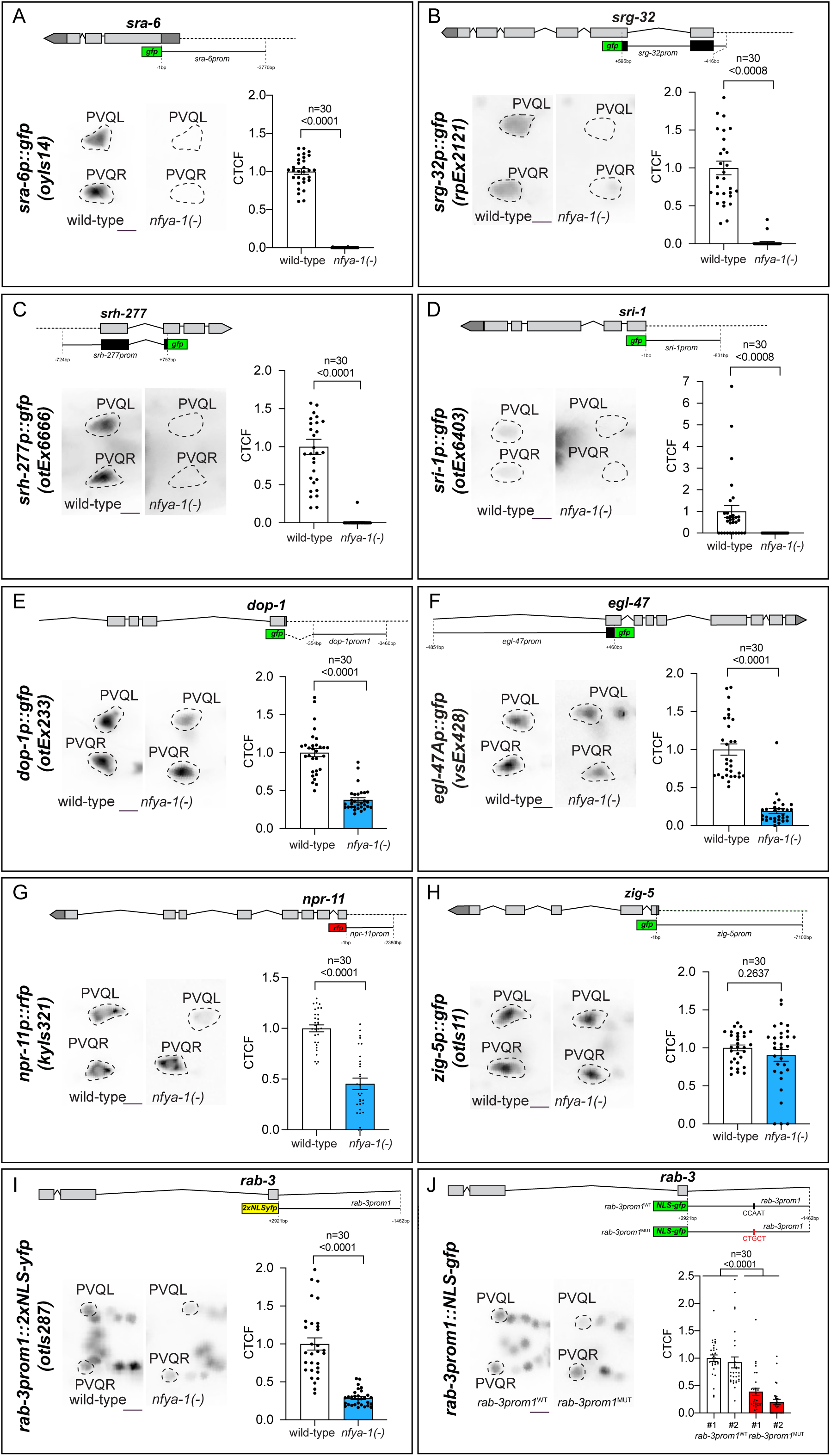
NFYA-1 controls neuron-type-specific and pan-neuronal expression in the PVQ interneurons. (A-I) Fluorescent micrographs and quantification of fluorescence intensity (CTCF= calculated total fluorescence) of neuron-type-specific (A-H) and pan-neuronal (I) reporter gene expression in the PVQs of wild-type and *nfya-1(ok1174)* animals. Schematics show the length in base pairs (bp) of each promoter in relation to the translational start site (ATG) for each reporter gene. n = 30 neurons per bar. Data expressed as mean ± SEM and statistical significance assessed by unpaired t test. Dashed circles show PVQ expression. Scale bar: 10 μm. (J) Fluorescent micrographs and quantification of fluorescence intensity (CTCF= calculated total fluorescence) of gene expression for wild-type and NFYA-1 binding site mutant (CCAAT>CTGCT; +1790-1974 bp) *rab-3prom1* reporters. n = 30 neurons per bar. Data expressed as mean ± SEM and statistical significance assessed by one-way ANOVA with Tukey’s correction. Dashed circles indicate PVQ expression. Scale bar: 10 μm.

We wondered whether NFYA-1 also controls pan-neuronal gene expression in the PVQ neurons. We examined the expression of a transcriptional reporter for the RAB-3 GTPase (*rab-3prom1::2xNLS-yfp*), a pan-neuronally-expressed protein that controls synaptic vesicle release (Nonet et al., 1997). We found that NFYA-1 is required for *rab-3prom1::2xNLS-YFP* expression in the PVQs (Figure 2I). The *rab-3* promoter contains a putative CCAAT NF-Y binding site, suggesting a direct regulatory relationship. We tested this hypothesis by generating wild-type and CCAAT-mutated *rab-3prom1::NLS::gfp* transgenes (Figure 2J). We found that mutation of the CCAAT motif reduced the expression of *rab-3prom1*-driven transgenes in the PVQ neurons (Figure 2J). Therefore, NFYA-1 is important for neuron-type specific and pan-neuronal gene expression in the PVQ neurons.

### NFYA-1 Binds to Pan-Neuronal Gene Promoters

Previous studies have shown that specific terminal selector TFs regulate neuron-type-specific gene expression, whereas CUT homeodomain proteins can control pan-neuronal expression independently and in a modular fashion with terminal selector TFs (Leyva-Diaz and Hobert, 2022; Stefanakis et al., 2015). Our finding that NFYA-1 is required for correct expression of both neuron-type-specific and pan-neuronal features in the PVQ neurons prompted us to investigate whether this role for NFYA-1 extends throughout the nervous system.

We examined independent embryonic and L3 larval whole-animal chromatin immunoprecipitation followed by sequencing (ChIP-Seq) profiles of GFP-tagged NFYA-1, performed by the modENCODE consortium, to determine NFYA-1 binding to neuronal promoters (Davis et al., 2018). Bioinformatic analysis of the ChIP-Seq data identified 3264 embryonic and 2693 larval NFYA-1 binding peaks, with an overlap of 2021 genes (82%), suggesting that NFYA-1 regulates similar gene sets across development (Table S2). Most NFYA-1 binding peaks (>90%) are located between 0 and 3 kb upstream of a transcription factor start site, suggesting that NFYA-1 functions at promoters and enhancers to control gene expression, as in other systems (Figure 3A-B) (Dolfini et al., 2009). The ChIP dataset contained two previously identified NFYA-1 target genes, *egl-5* and *tbx-2*, demonstrating the robustness of this analysis (Deng et al., 2007; Milton et al., 2013). Motif discovery using Homer (http://homer.ucsd.edu/homer/) identified a 10 bp sequence overrepresented in the NFYA-1 binding peaks that resembles the NFYA-1 binding site of mammalian orthologs (Figure 3C) (Dolfini et al., 2009). We were curious whether NFYA-1 binding peaks were found in neuron-type-specific and/or pan-neuronal genes. As NFYA-1 is broadly expressed, gene ontology (GO) analysis of genes containing NFYA-1 peaks did not identify overrepresented biological processes in the nervous system. Therefore, we took an unbiased approach, asking whether 15 randomly picked neuron-type-specific and pan-neuronal genes contain NFYA-1 ChIP peaks. We found that NFYA-1 binds to *cis*-regulatory elements of 1/15 neuron-type-specific genes (many of which are broadly expressed) and 15/15 pan-neuronal genes, suggesting that NFYA-1 predominantly controls the expression of pan-neuronal genes in *C. elegans* (Figure 3D-E).

**Figure 3.**
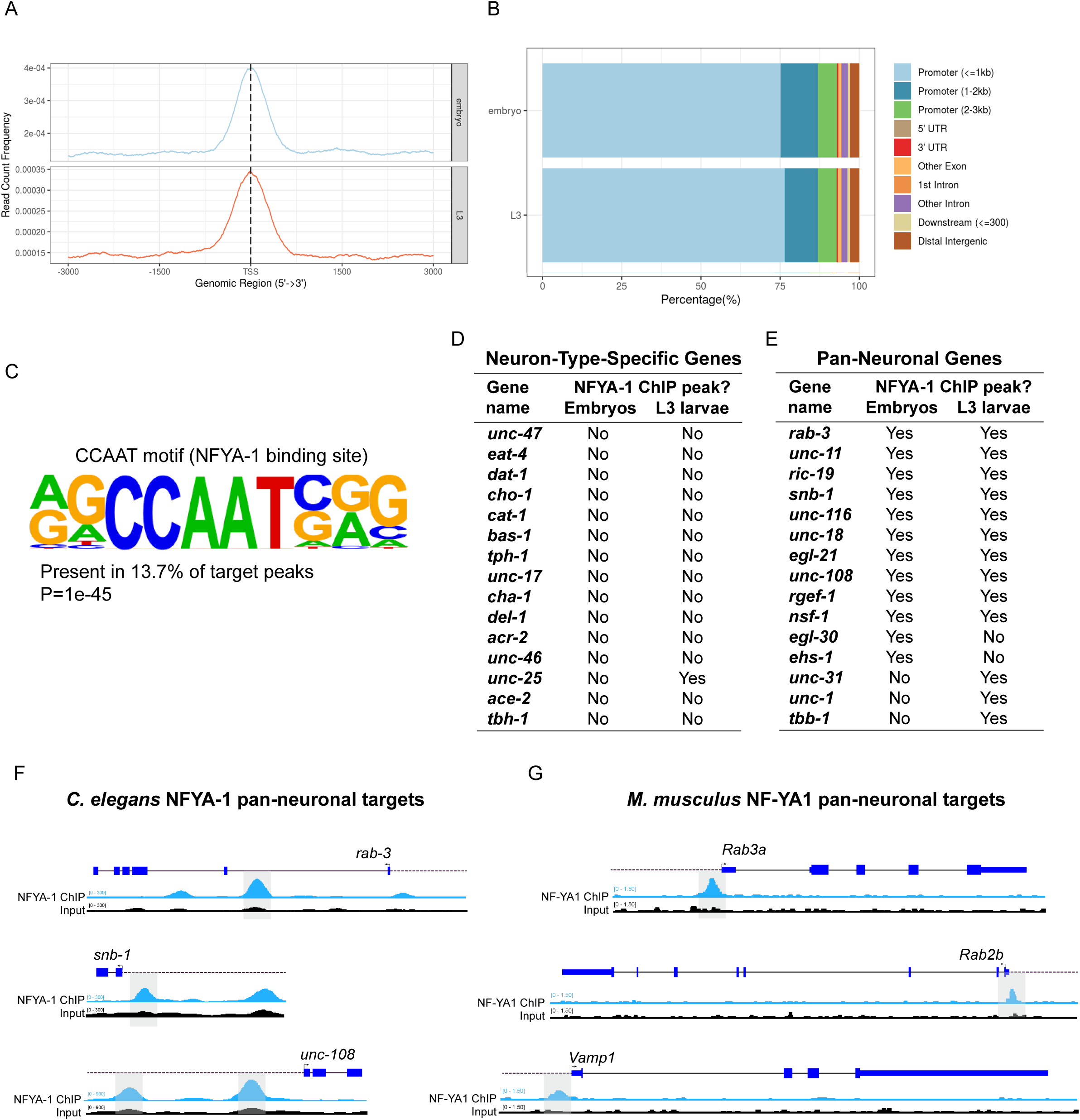
Mapping NFYA-1 genome-wide binding with ChIP-Sequencing. (A) Summary plot of NFYA-1 ChIP-Seq signals from embryos and L3 larvae (−3 kb and +3 kb from the TSS). (B) Summary of the NFYA-1 ChIP-Seq signal genomic distribution. (C) Motif discovery analysis of 3270 NFYA-1 binding peaks identified a 10 bp-long NFYA-1 binding motif. (D-E) Presence of NFYA-1 binding peaks in neuron-type-specific (D) and pan-neuronal (E) genes in embryos and L3 larvae. (F-G) Schematic representation of NFYA-1 ChIP-Seq (blue tracks) and input control (black tracks) in *C. elegans* (F) and *Mus musculus* (G). The *cis*-regulatory regions of three pan-neuronal genes in *C. elegans* (*rab-3*, *snb-1* and *unc-108*) and their mouse orthologs (*Rab3A*, *Rab2b* and *Vamp1*) show conserved NFYA-1 binding (highlighted by grey bars)

We surveyed publicly available ChIP-Seq datasets from terminally differentiated murine neurons (Zou et al., 2022) to determine whether NFYA-1 binding to pan-neuronal genes is conserved (Table S3). Of the 4085 genes bound by mouse NF-YA1 in neurons, 3040 genes have identifiable *C. elegans* orthologs (Table S3). We investigated whether any of the pan-neuronal genes bound by *C. elegans* NFYA-1 could also be controlled by murine *NF-YA1*. We found that three *C. elegans* pan-neuronal genes bound by NFYA-1 (*rab-3*, *unc-108* and *snb-1*) are also bound by *NF-YA1* (*Rab3a*, *Rab2b* and *Vamp1*) in mice (Figure 3F-G). These data suggest that NFYA-1 performs an evolutionarily conserved function in the control of pan-neuronal gene expression.

### NFYA-1 Controls Pan-Neuronal Gene Expression

We assessed the functional importance of NFYA-1 binding to pan-neuronal gene promoters by examining the expression of endogenous-, fosmid-, and promoter-based fluorescent reporters for genes that encode factors with general functions in synaptic transmission (*rab-3/RAB3, unc-11/SNAP91*, *ric-19/ICA1,* and *snb-1/RAB2B*) and neuropeptide processing (*egl-3/PCSK2*) (Figure 4 and S4). We measured fluorescence of *rab-3*, *unc-11*, *ric-19* and *egl-3* reporters in five (DB5, AS6, VD7, DA5 and VC5) and *snb-1* in four (VA6, VB7, DB5 and AS6) VNC motor neurons (Figure 4 and S4). These neurons represent distinct neuronal subclasses born at different developmental stages (Sulston, 1976). We detected reduced *rab-3* and *unc-11* expression, an increase in *snb-1* expression, and no change in *ric-19* or *egl-3* (Figure 4A-C and S4). These data reveal that NFYA-1 is important for the correct expression of specific pan-neuronal genes in *C. elegans* VNC motor neurons. However, the presence of an NFYA-1 binding peak is not sufficient for gene regulation, and thus additional regulatory factors likely cooperate with NFYA-1 to regulate neuronal gene expression in certain contexts (Figure S4).

**Figure 4.**
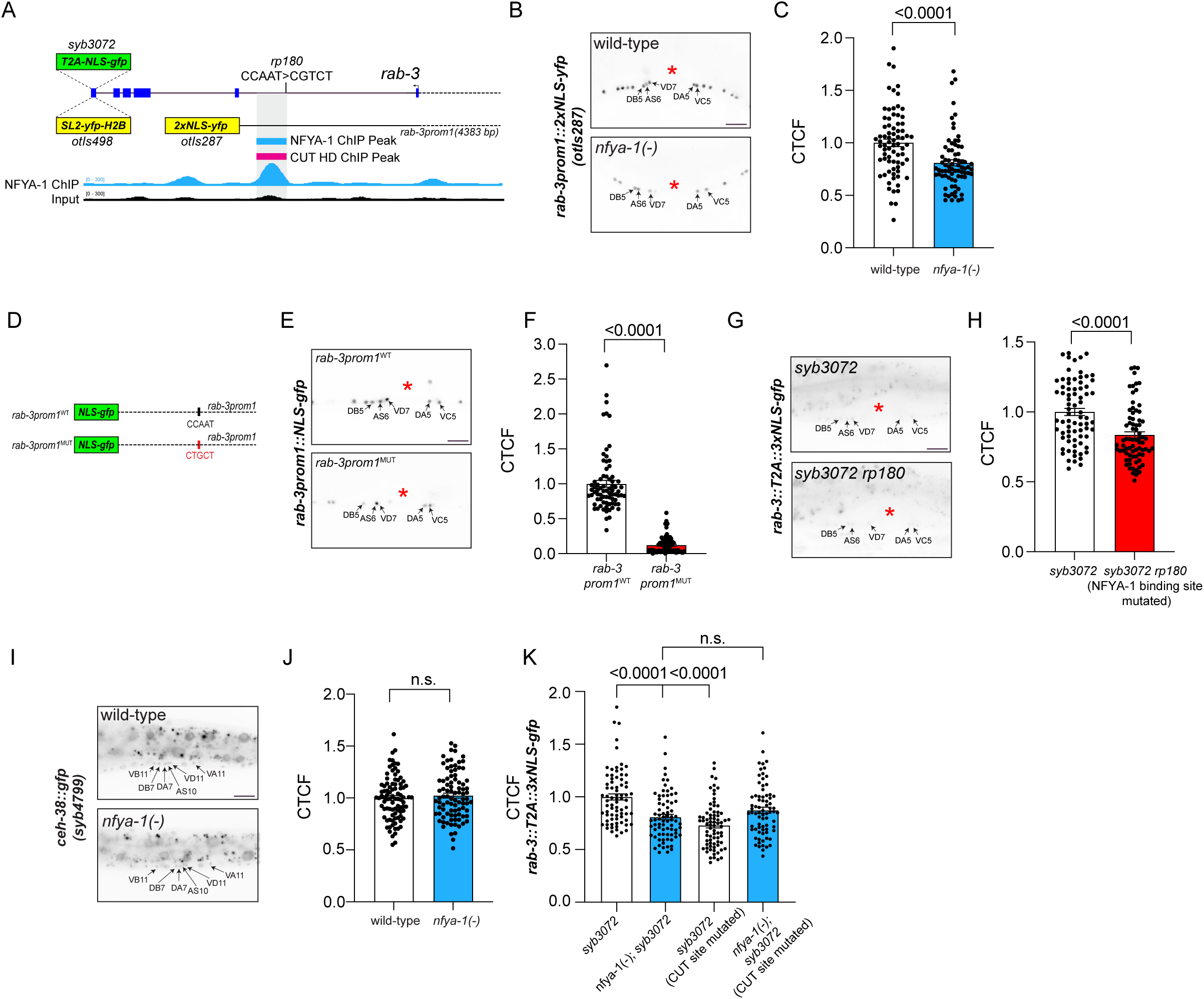
NFYA-1 controls pan-neuronal expression in motor neurons. (A) Schematic representation of the *rab-3* gene locus showing the location of the NFYA-1 ChIP peak (blue) and input control (black). Reporter genes used: *rab-3fosmid::SL2-yfp-H2B(otIs498*), *rab-3prom1::2xNLS-yfp(otIs287)* and endogenous *rab-3::T2A::3xNLS-gfp(syb3072)*. *rp180* = mutation of the putative NFYA-1 binding site (CCAAT>CGTCT) in *rab-3::T2A::3xNLS-gfp(syb3072)* animals. (B-C) Fluorescent micrographs (B) and quantification (C) of fluorescence intensity (CTCF= calculated total fluorescence) of a transcriptional *rab-3prom1::2xNLS-yfp(otIs287*) reporter in VNC motor neurons (DB5, AS6, VD7, DA5, and VC5) of wild-type and *nfya-1(ok1174)* animals. n = 75 neurons per bar. Data expressed as mean ± SEM and statistical significance assessed by unpaired t test. Red asterisk = vulva. Scale bar: 10 μm. (D-F) Schematic representation (D), fluorescent micrographs (E) and quantification (F) of fluorescence intensity (CTCF= calculated total fluorescence) of wild-type and NFYA-1 binding site mutant (CCAAT>CGTCT) *rab-3* transcriptional reporters in VNC motor neurons (DB5, AS6, VD7, DA5, and VC5) in wild-type animals. n = 75 neurons per bar. Data expressed as mean ± SEM and statistical significance assessed by unpaired t test. Red asterisk = vulva. Scale bar: 10 μm. (G-H) Fluorescent micrographs (G) and quantification (H) of fluorescence intensity (CTCF= calculated total fluorescence) of endogenously tagged *rab-3* in wild-type *rab-3::T2A::3xNLS-gfp(syb3072)* and upon mutation of the NFYA-1 binding site (CCAAT>CGTCT) *(rp180 syb3072[rab-3::T2A::3xNLS-gfp])* in the VNC motor neurons (DB5, AS6, VD7, DA5, and VC5) in wild-type animals. n = 75 neurons per bar. Data expressed as mean ± SEM and statistical significance assessed by unpaired t test. Red asterisk = vulva. Scale bar: 10 μm. (I-J) Fluorescent micrographs (I) and quantification (J) of fluorescence intensity (CTCF= calculated total fluorescence) of endogenously tagged *ceh-38::gfp(syb4799)* in VNC motor neurons (VB11, DB7, DA7, AS10, VD11 and VA11) of wild-type and *nfya-1(ok1174)* animals. n = 90 neurons per bar. Data expressed as mean ± SEM and statistical significance assessed by unpaired t test. Scale bar: 10 μm. (K) Quantification of fluorescence intensity (CTCF= calculated total fluorescence) of endogenously tagged *rab-3::T2A::3xNLS-gfp(syb3072)* in VNC motor neurons (DB5, AS6, VD7, DA5, and VC5) of wild-type and *nfya-1(ok1174)* animals with an intact or mutated CUT TF binding site. n = 75 neurons per bar. Data expressed as mean ± SEM and statistical significance was assessed by one-way ANOVA multiple comparison with Tukey’s correction. n.s., not significant.

We focused on *rab-3* regulation in VNC motor neurons to determine the functional relevance of putative NFYA-1 binding sites (Figure 4D-H). We mutated the NFYA-1 consensus sequence (CCAAT>CGTCT) that coincides with the NFYA-1 ChIP peak in the *rab-3* promoter (Figure 4A) and found that motor neuron expression was markedly reduced compared to the wild-type promoter-driven transgene (Figure 4D-F). Furthermore, introducing the same NFYA-1 binding site mutation using CRISPR-Cas9 editing in the *syb3072[rab-3::T2A::3xNLS::gfp])* endogenous reporter reduced expression in VNC motor neurons (Figure 4A and G-H). Therefore, NFYA-1 binding sites are required for faithful pan-neuronal gene expression.

### NFYA-1 Cooperates with Multiple TFs to Regulate Pan-Neuronal Expression

Loss of NFYA-1 does not eliminate pan-neuronal gene expression (Figure 4 and S4). A previous study similarly showed that six members of the CUT homeodomain TF family are redundantly required to partially control pan-neuronal gene expression (Leyva-Diaz and Hobert, 2022). We hypothesized that NFYA-1 controls pan-neuronal gene expression either by directly regulating CUT gene expression or by cooperating with CUT TFs at pan-neuronal gene loci. Analysis of ChIP-Seq data suggests that NFYA-1 does not directly control CUT gene expression (Table S2). This is supported by the finding that NFYA-1 loss has no effect on expression of the CUT gene *ceh-38::gfp(syb4799)* endogenous reporter (Figure 4I-J). As NFYA-1 and CUT TF ChIP binding peaks overlap at the *rab-3* promoter (Figure 4A), we investigated whether NFYA-1 and CUT TFs cooperate to control *rab-3* expression. We confirmed a previous report showing that endogenous *rab-3* expression is reduced when the CUT binding site, preventing any CUT TF binding, is mutated (Figure 4K) (Leyva-Diaz and Hobert, 2022). This CUT binding site mutation did not further reduce endogenous *rab-3* expression in *nfya-1(ok1174)* animals, suggesting that NFYA-1 and CUT TFs cooperatively control *rab-3* expression (Figure 4K). Whether NFYA-1 and CUT TFs act in a complex or that NFYA-1 is important for maintaining an optimal chromatin conformation for CUT TF binding to the *rab-3* promoter is an open question.

We were curious whether NFYA-1 can collaborate with other TFs to regulate pan-neuronal expression. The UNC-30/Pitx and UNC-55/COUP TFs are key regulators of neuron-type specific gene expression, but not pan-neuronal expression, in GABAergic neurons (Jin et al., 1994; Petersen et al., 2011; Stefanakis et al., 2015; Yu et al., 2017). We measured *rab-3fosmid::SL2-yfp-H2B* levels in the VD7 GABAergic motor neuron that co-expresses NFYA-1, UNC-30 and UNC-55 (Figure 5A-E). As an internal control, we also measured pan-neuronal expression in the neighboring cholinergic DB5 and AS6 neurons (Figure 5C-E). We generated single and double mutant loss-of-function combinations between *nfya-1(ok1174)*, *unc-30(e191)*, and *unc-55(e1170)* and discovered that removal of UNC-30 enhanced, and UNC-55 reversed the reduction of *rab-3* expression in the VD7 neuron of *nfya-1(ok1174)* animals (Figure 5C-E). In contrast, loss of UNC-30 and UNC-55 did not affect *rab-3* expression in the DB5 and AS6 neurons (Figure 5C-E). In addition, NFYA-1 loss reduces UNC-30 expression in VD7 (Figure 5F), suggesting combinatorial regulation of *rab-3* expression (Figure 5G). Together, these data reveal that NFYA-1 collaborates with multiple TFs to regulate pan-neuronal expression in specific contexts.

**Figure 5.**
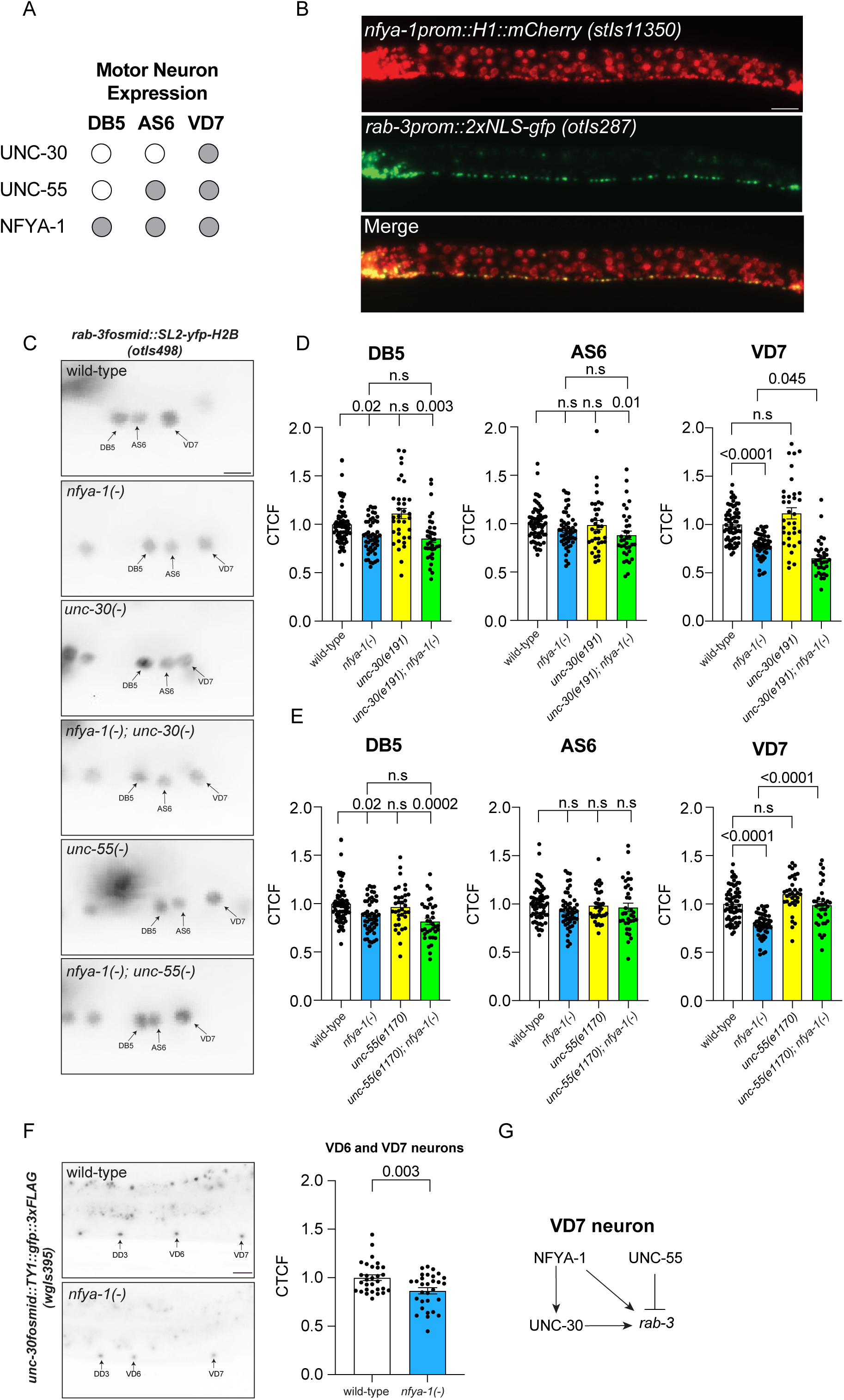
NFYA-1 cooperates with UNC-30/Pitx and UNC-55/COUP to control pan-neuronal gene expression in a GABAergic motor neuron. (A) Schematic representation expression of NFYA-1, UNC-30 and UNC-55 in the DB5, AS6, and VD7 VNC motor neurons. Grey = expressed, white = not expressed. (B) Fluorescent micrographs of *nfya-1prom::H1::mCherry (stIs11350)* (top), *rab-3prom::2xNLS-gfp (otIs287)* (middle), and merge (bottom). Scale bar = 10 μm. (C-E) Fluorescent micrographs (C) and quantification (D-E) of fluorescence intensity (CTCF= calculated total fluorescence) of endogenously tagged *rab-3::T2A::3xNLS-gfp(syb3072)* in VNC motor neurons (DB5, AS6 and VD7) of wild-type, *nfya-1(ok1174)*, *unc-30(e191)*, *nfya-1(ok1174); unc-30(e191)*, *unc-55(e1170)* and *nfya-1(ok1174); unc-55(e1170)* animals. n = >35 neurons per bar. Data expressed as mean ± SEM and statistical significance assessed by unpaired t test. Scale bar: 10 μm. (F) Fluorescent micrographs and quantification of fluorescence intensity (CTCF= calculated total fluorescence) of *unc-30fosmid::TY1::gfp::3xFLAG(wgIs395)* VD6-7 motor neurons. n = 30 neurons per bar. Data expressed as mean ± SEM and statistical significance assessed by unpaired t test. Scale bar: 10 μm. (G) Schematic model of the gene network that regulates *rab-3* expression in the VD7 GABAergic motor neuron.

### NFYA-1 Controls Neuron-Type-Specific Gene Expression

Neuron-type-specific genes, such as those encoding neurotransmitter biosynthesis and transport proteins, are regulated by terminal selector TFs, but not by CUT homeodomain proteins (Flames and Hobert, 2009; Gendrel et al., 2016; Leyva-Diaz and Hobert, 2022; Serrano-Saiz et al., 2013). Our analysis of ChIP-Seq data detected minimal NFYA-1 binding to neuron-type-specific gene promoters (Figure 3D and Table S2). However, we did observe NFYA-1-dependent regulation of some neuron-type-specific genes in certain neuronal contexts (Figure 6). We found that NFYA-1 loss causes downregulation of *eat-4/VGLUT* (a glutamatergic neuron-specific marker) in the URYD, OLL, IL1, OLQV, and BAG head neurons and upregulation of the following neuron-type-specific gene reporters: *unc-25/GAD* and *unc-47/VGAT* (GABAergic neuron-specific markers) in the DD neurons and *unc-47/VGAT* in the VD neurons, and *dat-1/DAT* (a dopaminergic neuron-specific marker) in the CEPD neurons (Figure 6A-D and S5A-C). In contrast, NFYA-1 loss did not detectably change expression of three cholinergic neuron-type-specific gene reporters we analyzed: *acr-2/AChR, cho-1/ChT* and *unc-17/ChAT* (Figure 6E-F and S5D-E). Thus, NFYA-1 impacts the expression of neuron-type-specific genes in a context-dependent and likely indirect manner (Figure 6G).

**Figure 6.**
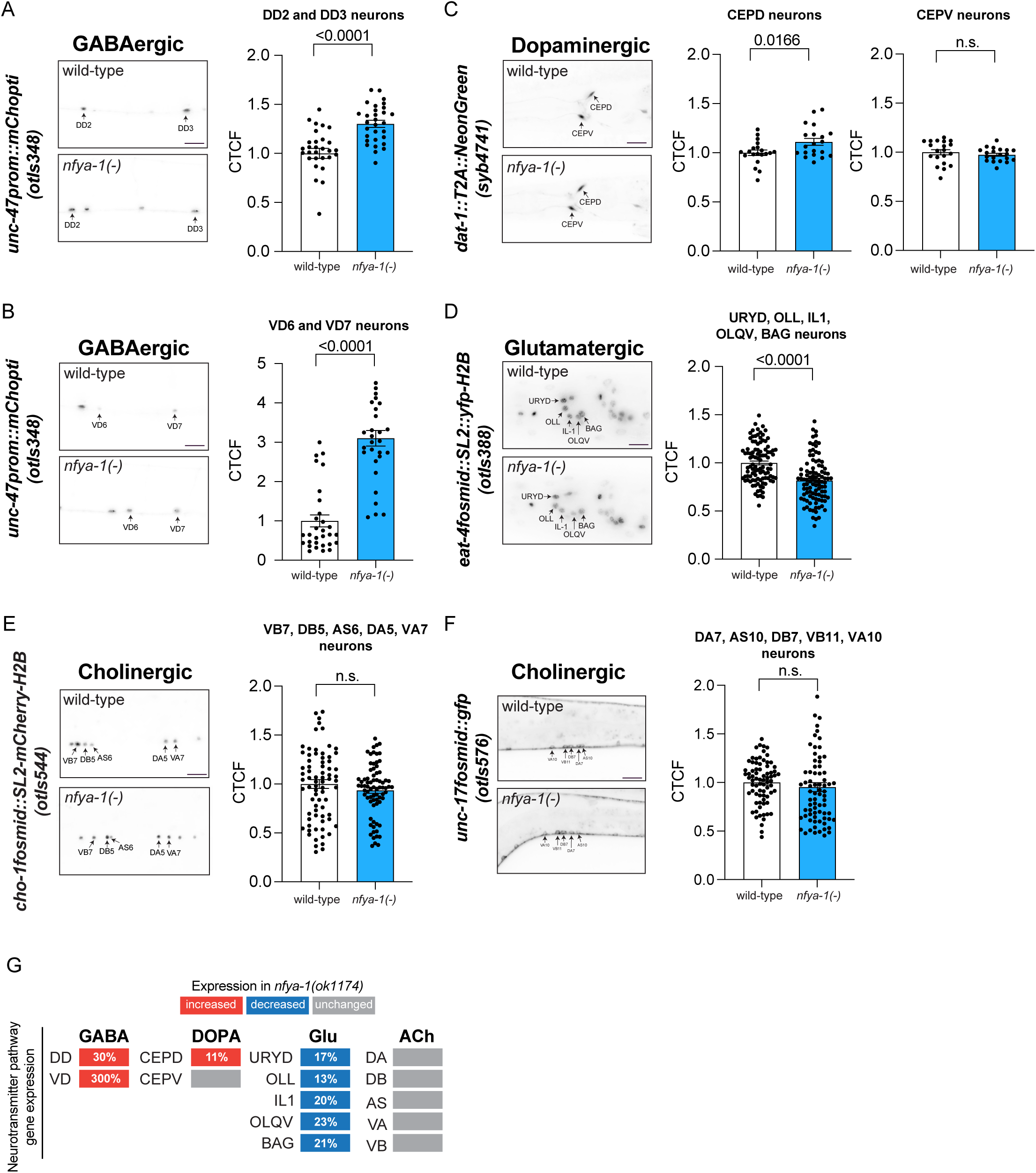
NFYA-1 regulation of neuron-type specific gene expression. (A-H) Fluorescent micrographs and quantification of fluorescence intensity (CTCF= calculated total fluorescence) of *unc-47prom::mChopti(otIs348)* (A-B), *dat-1::T2A::NeonGreen(syb4741)* (C), *eat-4fosmid::SL2::yfp-H2B(otIs388)* (D), *cho-1fosmid::SL2::mCherry-H2B(otIs544)* (E), and *unc-17fosmid::gfp(otIs576)* (F) in wild-type and *nfya-1(ok1174)* animals. n = >20 neurons per bar. Data expressed as mean ± SEM and statistical significance assessed by unpaired t test. Scale bars: 10 μm. (G) Effect of NFYA-1 loss on neuron-type specific gene expression. Colored rectangles denote expression increase (red), decrease (blue) or unchanged (grey) in *nfya-1(ok1174)* animals compared to wild-type. Percentage change shown for each individual neuron. GABA = GABAergic, DOPA = Dopaminergic, Glu = Glutamatergic and Ach = Cholinergic.

To investigate how NFYA-1 regulates neuron-type-specific gene expression, we focused on the *del-1* gene, which encodes a DEG/ENaC ion channel expressed in the VA/VB motor neurons (Tavernarakis et al., 1997). The *del-1::gfp* transcriptional reporter is expressed in the VB motor neurons from the early L2 larval stage and is weakly expressed in the VA motor neurons of L4 larvae (Figure 7A) (Winnier et al., 1999). We observed elevated *del-1::gfp* expression in the VA, but not VB, motor neurons in *nfya-1(ok1174)* animals at the L4 larval stage (Figure 7A-C). As no NFYA-1 ChIP peaks were detected at the *del-1* locus (Table S2), we hypothesized that NFYA-1 indirectly regulates *del-1::gfp* expression. The UNC-3/COE terminal selector TF is essential for *del-1::gfp* expression in both the VA/VB motor neurons (Figure 7D) (Kerk et al., 2017; Kratsios et al., 2017; Prasad et al., 2008). In addition, *del-1::gfp* expression is spatiotemporally regulated by the Prd-type homeobox protein UNC-4 and the Groucho-like co-repressor UNC-37 (Winnier et al., 1999). UNC-4 and UNC-37 prevent precocious *del-1::gfp* expression in the VA motor neurons, which likely enables correct synaptic wiring (Figure 7D) (Winnier et al., 1999). We wondered whether *del-1::gfp* derepression in *nfya-1(ok1174)* animals is caused by dysregulation of one or all of UNC-3, UNC-4, and UNC-37. We found that NFYA-1 is required for complete expression of endogenous reporters for UNC-3, UNC-4, and UNC-37 in VNC motor neurons, including the VAs (Figure 7E-G). Unfortunately, we were unable to generate compound mutants between *nfya-1(ok1174)* and null mutants for *unc-3* and *unc-37* due to lethality. However, we were able to combine *nfya-1(ok1174)* with the *unc-4(e120)* presumptive null allele. We measured *del-1::gfp* expression in the VA neurons of *nfya-1(ok1174); unc-4(e120)* animals and did not detect an additive effect (Figure 7H-I). In fact, we detected lower *del-1::gfp* expression than the *unc-4(e120)* single mutant (Figure 7H-I). We suggest that the reduction in *del-1::gfp* expression observed in *nfya-1(ok1174); unc-4(e120)* animals compared to *unc-4(e120)* is due to lower UNC-3 expression, and thus reduced transcriptional activation by UNC-3 at the *del-1* promoter (Figure 7J).

**Figure 7.**
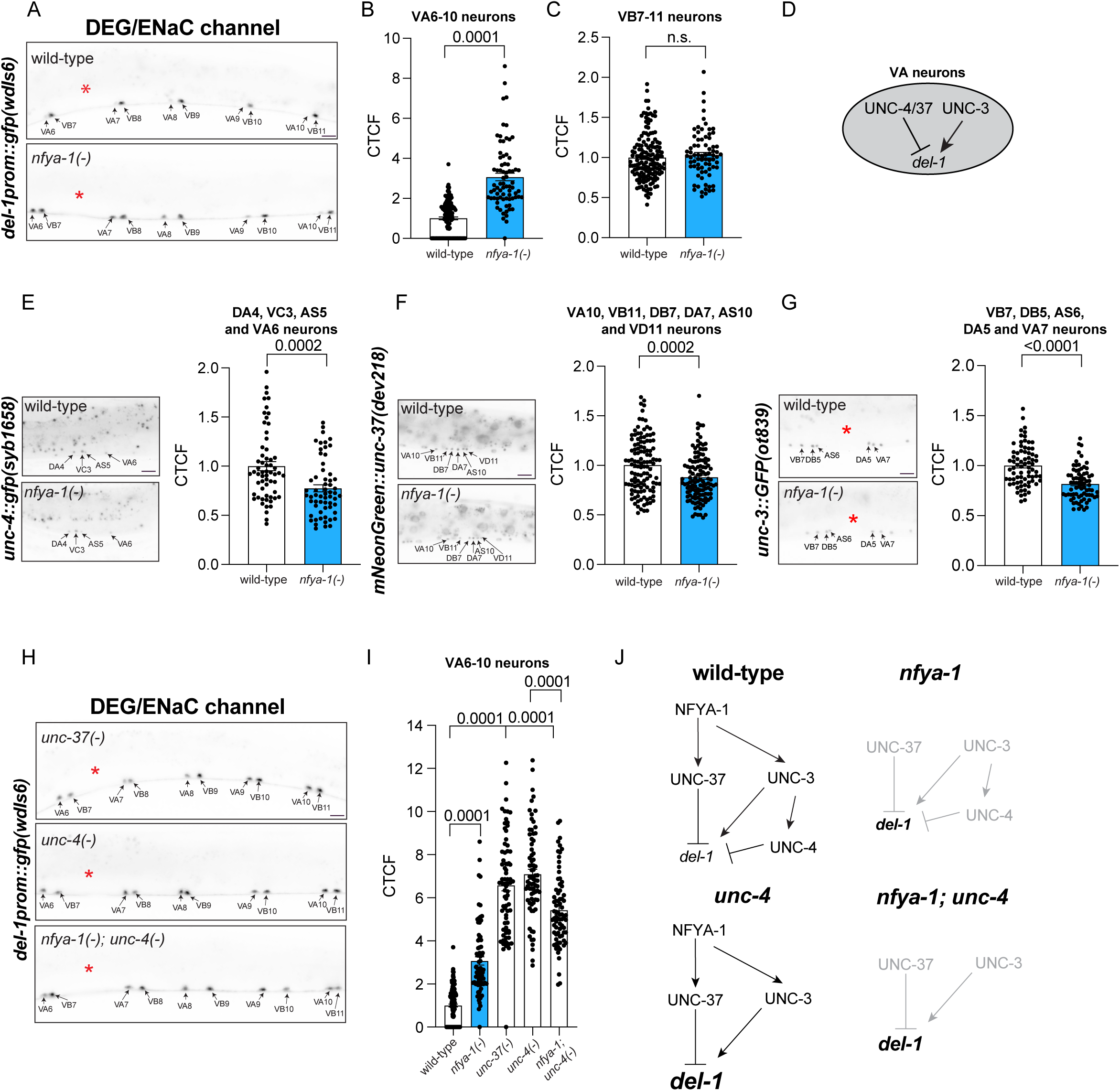
NFYA-1 regulates three TFs that coordinate degenerin channel expression. (A-C) Fluorescent micrographs (A) and quantification (B-C) of fluorescence intensity (CTCF= calculated total fluorescence) of *del-1prom::gfp(wdIs6)* in VNC motor neurons (VA6-10 (B) and VB7-11 (C)) of wild-type and *nfya-1(ok1174)* animals. n >75 neurons per bar. Data expressed as mean ± SEM and statistical significance assessed by unpaired t test. Red asterisk = vulva. n.s., not significant. Scale bar: 10 μm. (D) Mechanism for regulating *del-1prom::gfp* expression in the VA neurons. (E-G) Fluorescent micrographs and quantification of fluorescence intensity (CTCF= calculated total fluorescence) of endogenous reporters (*unc-4::gfp(syb1658)* (E), *mNeonGreen::unc-37(dev218)* (F), and *unc-3::gfp(ot839)* (G) in wild-type and *nfya-1(ok1174)* animals. n >60 neurons per bar. Data expressed as mean ± SEM and statistical significance assessed by unpaired t test. Red asterisk = vulva. Scale bar: 10 μm. (H-I) Fluorescent micrographs (H) and quantification (I) of fluorescence intensity (CTCF= calculated total fluorescence) of *del-1prom::gfp(wdIs6)* in VNC motor neurons (VA6-10 of wild-type, *nfya-1(ok1174)*, *unc-37(e262)*, *unc-4(e120)*, and *nfya-1(ok1174)*; *unc-4(e120)* animals. (Note - wild-type and *nfya-1(ok1174)* micrographs are shown in panel A). n >75 neurons per bar. Data expressed as mean ± SEM and statistical significance was assessed by one-way ANOVA multiple comparison with Tukey’s correction. Red asterisk = vulva. n.s., not significant. Scale bar: 10 μm. (J) Illustration of how NFYA-1 controls *del-1* expression through the regulation of multiple transcription factors.

### The NF-Y Complex Controls Neuronal Activity and Behavior

Our data show that NFYA-1 regulates genes responsible for neurotransmitter generation and transport (*unc-25*/*GAD*, *unc-47/VGAT, eat-4/VGLUT*, and *dat-1/DAT*) and synaptic function (*rab-3/RAB3, unc-11/SNAP91*, and *snb-1/RAB2B*) (Lee et al., 1999; McIntire et al., 1993; Nass et al., 2002; Nonet et al., 1999; Nonet et al., 1998; Nonet et al., 1997). While, in most cases NFYA-1 loss caused moderate changes in neuronal gene expression, we wondered whether their cumulative effect would affect neuronal activity and/or animal behavior. Therefore, we measured the motility of *C. elegans* adult hermaphrodites in liquid medium using the thrash assay (Miller et al., 1996). We found that one-day old *nfya-1(ok1174)* animals exhibited fewer thrashes per minute than wild-type animals, and this phenotype was rescued by re-supplying a *nfya-1*-containing fosmid transgene (Figure 8A). Next, we examined the locomotory speed (distance travelled/time) of one-day old *nfya-1(ok1174)* and *nfya-1(bp4)* adult hermaphrodites using WormLab automated tracking (MBF Bioscience). Each *nfya-1* mutant allele exhibited reduced locomotion compared to wild-type (Figure 8B). We rescued the *nfya-1(ok1174)* motility defect by driving *nfya-1* cDNA pan-neuronally (Figure 8C). Furthermore, simultaneous expression of *nfya-1* cDNA in cholinergic and GABAergic motor neurons, using the *unc-3* and *unc-47* promoters, also partially rescued the locomotion defects of *nfya-1(ok1174)* animals (Figure 8D). The locomotion defects observed in *nfya-1(ok1174)* animals may be caused, at least in part, by disrupted VNC motor neuron activity. We therefore expressed a chameleon ratiometric calcium sensor (YC3.60) in the nervous system and used Förster resonance energy transfer (FRET) microscopy to measure calcium levels in VNC motor neurons of immobilized animals. We detected a small but statistically significant reduction in neuronal activity in motor neurons of *nfya-1(ok1174)* animals (Figure 8E-F). Together, these data suggest that the cumulative effect of disrupted neuronal gene expression in VNC motor neurons when NFYA-1 is absent is defective motor neuron activity and function.

**Figure 8.**
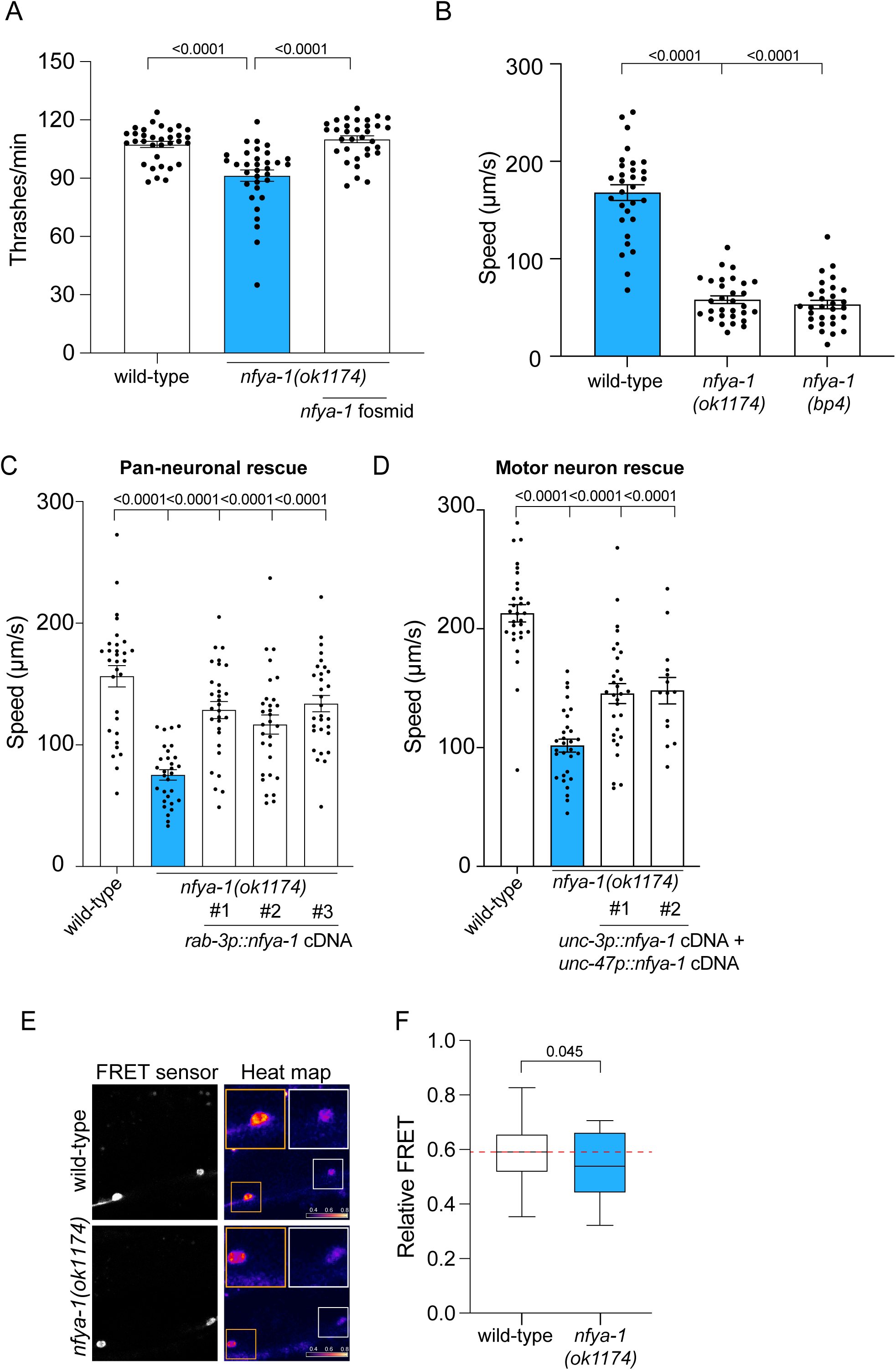
NFYA-1 controls neuronal activity and animal behavior. (A) Number of thrashes per minute in wild-type and *nfya-1(ok1174)* animals. Expressing an *nfya-1*-expressing fosmid (WRM0633cC02) restores wild-type thrashing to *nfya-1(ok1174)* mutant animals. n = 30 per bar. Data expressed as mean ± SEM and statistical significance assessed by one-way ANOVA with Dunnet’s correction. (B) Quantification of locomotory speed (distance travelled/time) in wild-type, *nfya-1(ok1174)*, and *nfya-1(bp4)* animals. n = 33 per bar. Data expressed as mean ± SEM and statistical significance was assessed by one-way ANOVA multiple comparison with Tukey’s correction. (C-D) Quantification of locomotory speed (distance travelled/time) of wild-type, *nfya-1(ok1174),* and *nfya-1(ok1174)* mutant rescue (pan-neuronal (C) and motor neuron (D) promoters). n = 30 per bar. Data expressed as mean ± SEM and statistical significance was assessed by one-way ANOVA multiple comparison with Tukey’s correction. (E-F) Fluorescent micrographs (E) and quantification (F) of calcium imaging by FRET microscopy of VNC motor neurons from live immobilized wild-type and *nfya-1(ok1174)* animals expressing a chameleon sensor, YC3.60. n = 67-79. Data expressed as mean ± SEM and statistical significance assessed by unpaired t test with Welch’s correction.

## DISCUSSION

Here, we have shown that the NFYA-1 TF is a key regulator of neuron-type-specific and pan-neuronal gene expression. Previous research demonstrated that neuron-type-specific (ion channels, neurotransmitter biosynthetic enzymes, and neuropeptide receptors) and pan-neuronal (neuropeptide processing and synaptic vesicle release) gene batteries are regulated through distinct mechanisms. While neuron-type-specific gene expression is controlled by terminal selector TFs, pan-neuronal gene expression is regulated by multiple redundant TFs. For example, the UNC-3/COE and UNC-30/Pitx terminal selector TFs are necessary for the expression of cholinergic and GABAergic neuron-type-specific genes, respectively (Eastman et al., 1999; Jin et al., 1994; Kratsios et al., 2011). Yet these TFs are expendable for pan-neuronal gene expression in these neuron subclasses (Kratsios et al., 2011; Leyva-Diaz and Hobert, 2022). In contrast, the CUT homeodomain TF family controls pan-neuronal gene expression directly and in a cooperative manner with terminal selector TFs, but does not control neuron-type-specific gene expression (Leyva-Diaz and Hobert, 2022). The redundant inputs that regulate pan-neuronal genes are thought to provide robustness to these critical gene batteries that control general neuronal functions (Leyva-Diaz and Hobert, 2022). Our findings reveal an additional layer of neuronal gene regulation governed by the NFYA-1 TF, which, unlike previously identified factors, regulates both neuron-type-specific and pan-neuronal genes.

We discovered a role for the NF-Y TF complex in the nervous system through an unbiased genetic screen for regulators of PVQ interneuron fate. We found that NFYA-1 is required for the expression of many neuron-type-specific genes in the PVQ neurons, as well as the requirement for NF-Y binding sites for PVQ expression - indicating a direct mode of regulation. Furthermore, we demonstrated that NFYA-1 acts cell autonomously and likely from early development to control PVQ fate. Surprisingly, we revealed that NFYA-1 loss also decreased expression of a pan-neuronal reporter gene (*rab-3/RAB3*) in the PVQ neurons, and that mutating the NF-Y binding site in the *rab-3* promoter lowered expression. This finding prompted us to investigate whether NFYA-1 controls neuronal gene expression more broadly.

Our analysis of NFYA-1 ChIP-Seq data revealed prominent binding to the promoters of pan-neuronal genes, with little binding to neuron-type-specific gene promoters. Our examination of NF-YA1 ChIP-Seq data from murine neurons revealed an analogous theme, suggesting conserved NF-Y complex binding and regulation of neuronal gene batteries. As predicted by the presence of NFYA-1 ChIP peaks, we found that the expression of multiple pan-neuronal genes is dependent on NFYA-1. Focusing on expression in VNC motor neurons, we found that NFYA-1 regulation of the pan-neuronal gene *rab-3* is dependent on a CCAAT NF-Y binding site that overlaps with the NFYA-1 ChIP peak. This pan-neuronal regulatory function parallels that of the CUT homeodomain TF family (Leyva-Diaz and Hobert, 2022), and we found that NFYA-1 likely collaborates with the CUT TFs to control the expression of specific pan-neuronal genes.

Surprisingly, despite the lack of NFYA-1 ChIP peaks, certain neuron-type-specific genes are also dysregulated in animals lacking NFYA-1. These include genes involved in GABAergic and dopaminergic synthesis and transport (*unc-25/GAD, unc-47/VGAT*, and *dat-1/DAT*) and the DEG/ENaC ion channel, DEL-1. We also observed distinct effects of NFYA-1 loss on the same neuron-type-specific reporter gene in different neuronal subclasses. For example, *unc-25/GAD* expression is increased in embryonic DD but not in post-embryonic VD GABAergic neurons. This may indicate temporally distinct regulation of TF expression and/or binding by NFYA-1 to specific promoter elements. Together, these changes in neuron-type-specific expression are unlikely to be a general response to defective pan-neuronal gene expression as: 1) these effects are not observed when CUT TF function is lost, and 2) the expression of other neuron-type-specific genes such as *cho-1/ChT* and *egl-3/PCSK2* were unaffected by NFYA-1 loss.

We found that NFYA-1 loss caused gene expression to be abolished in certain neurons, such as the PVQs, as also shown recently for the IL1 neurons. (Heo et al., 2022). However, the majority of neuronal genes analyzed showed ∼20-40% changes in gene expression. These subtler gene expression changes likely combine to cause defects in neuronal activity and locomotion observed in *nfya-1* knockout animals. Interestingly, a discordant twin study showed that hypermethylation of the human *NF-YC* promoter is observed in individuals with autism spectrum disorder (Wong et al., 2014). Thus, repressed NFY transcriptional activity, and the likely manifold associated changes in neuronal gene expression, may account for altered brain function in these individuals.

## SUPPLEMENTARY MATERIAL

### SUPPLEMENTARY FIGURES

**Figure S1. NF-Y complex gene structures and mutations in *C. elegans***

Schematics of gene structures for *nfya-1*, *nfya-2*, *nfyb-1* and *nfyc-1*. Exons = light grey boxes, untranslated regions = dark grey boxes, introns = black lines, genetic lesions = red lines.

**Figure S2. NFYA-1 Acts Cell Autonomously to Control PVQ Interneuron Fate**

(A) Fluorescent micrographs of *nfya-1prom::H1::mCherry*, *nfyb-1prom::H1::mCherry* and *nfyc-1prom::H1::mCherry* expression in the tail (left), with *sra-6prom::gfp(oyIs14)* in the PVQ neurons (centre) and merge (right). Dashed circles show PVQ expression. Scale bar = 10 μm.

(B) Quantification of *sra-6prom::gfp(oyIs14)* expression in the PVQ interneurons of wild-type and *nfya-1(ok1174)* animals. Expressing a *nfya-1*-expressing fosmid (WRM0633cC02) or PVQ-specifically in the PVQ neurons (*npr-11prom::nfya-1* cDNA) restores PVQ expression to *nfya-1(ok1174)* mutant animals. Data expressed as mean ± SEM and statistical significance assessed by one-way ANOVA with Dunnet’s correction. # = independent transgenic lines.

(C) Quantification of *sra-6prom::gfp(oyIs14)* expression in the PVQ interneurons of wild-type and *nfya-1(ok1174)* animals at the L1 and L4 stages of larval development. Data expressed as mean ± SEM and statistical significance assessed by one-way ANOVA with Dunnet’s correction. n.s., not significant.

**Figure S3. Putative NF-Y binding sites are required for PVQ expression**

(A) The number of putative NF-Y binding sites (CCAAT or ATTGG) located in PVQ expressed promoters. The length of each promoter used to drive fluorescent proteins is shown in bp.

(B-C) Quantification of reporter expression of *sri-1* (B) and *srh-277* (C) wild type and CCAAT binding site mutated promoters. Schematics shows the length in bp of each promoter in relation to the translational start site (ATG) for each reporter gene. The position of the mutated NF-Y binding site (CCAAT>CTGCT) binding site to the ATG is marked in red.

Data expressed as mean ± SEM and statistical significance assessed by one-way ANOVA with Tukey’s correction. n.s., not significant. # = independent transgenic lines.

**Figure S4. NFYA-1 regulation of pan-neuronal gene expression**

(A) Schematic representation of the *unc-11* gene locus showing the location of NFYA-1 ChIP peak (red) and input control (black). Reporter gene used: *unc-11fosmid::SL2-yfp-H2B(rpEx2212*).

(B-C) Fluorescent micrographs (B) and quantification (C) of fluorescence intensity (CTCF= calculated total fluorescence) of the *unc-11fosmid::SL2-yfp-H2B(rpEx2212*) reporter in VNC motor neurons (DB5, AS6, VD7, DA5, and VC5) of wild-type and *nfya-1(ok1174)* animals. n = 75 neurons per bar. Data expressed as mean ± SEM and statistical significance assessed by unpaired t test. Red asterisk = vulva. Scale bar = 10 μm.

(D) Schematic representation of the *snb-1* gene locus showing the location of NFYA-1 ChIP peak (red) and input control (black). Reporter gene used: *snb-1fosmid::SL2-yfp-H2B(rpEx2211*).

(E-F) Fluorescent micrographs (E) and quantification (F) of fluorescence intensity (CTCF= calculated total fluorescence) of the *snb-1fosmid::SL2-yfp-H2B(rpEx2211*) reporter in VNC motor neurons (VA6, VB7, DB5 and AS6) of wild-type and *nfya-1(ok1174)* animals. n = 75 neurons per bar. Data expressed as mean ± SEM and statistical significance assessed by unpaired t test. Red asterisk = vulva. Scale bar = 10 μm. Scale bar = 10 μm.

(G) Schematic representation of the *ric-19* gene locus showing the location of NFYA-1 ChIP peak (red) and input control (black). Reporter gene used: *ric-19prom::ric-19::gfp(etIs1*).

(H-I) Fluorescent micrographs and quantification of fluorescence intensity (CTCF= calculated total fluorescence) of the *ric-19prom::ric-19::gfp(etIs1*) reporter in VNC motor neurons (DB5, AS6, VD7, DA5, and VC5) of wild-type and *nfya-1(ok1174)* animals. n = 75 neurons per bar. Data expressed as mean ± SEM and statistical significance assessed by unpaired t test. Red asterisk = vulva. Scale bar = 10 μm. Scale bar = 10 μm.

(J) Schematic representation of the *egl-3* gene locus showing the location of NFYA-1 ChIP peak (red) and input control (black). Reporter gene used: *egl-3::SL2-gfp-H2B(syb4478)*.

(K-L) Fluorescent micrographs (K) and quantification (L) of fluorescence intensity (CTCF= calculated total fluorescence) of the endogenous *egl-3::SL2-gfp-H2B(syb4478)* reporter in VNC motor neurons (DB5, AS6, VD7, DA5, and VC5) of wild-type and *nfya-1(ok1174)* animals. n = 75 neurons per bar. Data expressed as mean ± SEM and statistical significance assessed by unpaired t test. Red asterisk = vulva. Scale bar = 10 μm.

**Figure S5. NFYA-1 Regulates Certain Neuron-Type-Specific Genes**

(A-H) Fluorescent micrographs and quantification of fluorescence intensity (CTCF= calculated total fluorescence) of *unc-25prom::gfp(juIs76)* (A-B), *dat-1prom::gfp(vtIs1)* (C), *unc-17prom::gfp(vsIs48)* (D) and *acr-2prom::gfp(juIs14)* (E). n >20 neurons per bar. Data expressed as mean ± SEM and statistical significance assessed by unpaired t test. Scale bars: 10 μm.

### SUPPLEMENTARY TABLES

**Table S1. Source data**

**Table S2. NFYA-1 target genes (C. elegans ChIP-Sequencing analysis)**

**Table S3. NF-YA1 target genes (*M. musculus* ChIP-Sequencing analysis)**

## EXPERIMENTAL PROCEDURES

### EXPERIMENTAL MODEL AND SUBJECT DETAILS

#### *C. elegans* strains and genetics

All *C. elegans* strains were maintained at 20°C on NGM plates seeded with *Escherichia coli* OP50 bacteria. Strains were generated using standard genetic procedures and are listed in the Key Resources Table. All strains were backcrossed to N2 at least three times before scoring or generating compound mutants. Genotypes were confirmed using PCR genotyping or Sanger sequencing with primers listed in the Key Resources Table.

#### Neuroanatomical observation and scoring

Neurons were scored blinded to genotype. The developmental stage and scoring criteria are detailed in each Figure legend. Transgenes used to visualize specific neurons are detailed in the Key Resources Table. All neuronal scoring was repeated in triplicate on independent days.

#### Forward mutagenesis screen

Ethyl methanesulfonate (EMS) mutagenesis was performed as previously described (Flibotte et al., 2010). F2 progeny of EMS-mutagenized P0 hermaphrodites carrying the *sra-6prom::gfp(oyIs14)* transgene were screened for abnormal PVQ expression under a dissecting microscope with a fluorescence light source.

#### Genetic mapping of *nfyc-1(rp120)*

We used the one-step whole-genome sequencing and SNP mapping strategy to map the *rp120* genetic lesion (Doitsidou et al., 2010). Males of the Hawaiian strain CB4856 were crossed with *rp120; oyIs14* hermaphrodites. Ten F1 progeny carrying the *oyIs14* transgene were picked to individual plates and allowed to self-fertilize. 50 F2 progeny carrying the *oyIs14* transgene with weak/loss of PVQ expression were picked and allowed to self-fertilize. Progeny of these animals were pooled and DNA isolated. Pooled genomic DNA was sequenced using Ilumina sequencing and the resultant sequencing data was analyzed using the Galaxy platform. Our mapping identified a single lesion (CAG-CAA) at the splice acceptor of exon 4 of the *nfyc-1* gene, which was independently confirmed by Sanger sequencing.

#### Microinjection and transgenic animal generation

Rescue and overexpression plasmids were microinjected into wild-type or mutant hermaphrodites at 1-50 ng/μl concentration with *PvuII* digested bacterial DNA (100-150 ng/μl) and *myo-2prom::mCherry* (1-5 ng/μl) as an injection marker.

#### Molecular cloning

Cloning of inserts was performed using a traditional restriction endonuclease or restriction-free protocols. Amplified inserts and plasmid backbones were digested using specific restriction enzymes, shown in the Key Resources Table. A single restriction enzyme reaction (50 μl) consisted of 600 ng of DNA, 1 μl of Fast Digest (Thermo scientific) restriction enzyme, 5 μl of 10X FastDigest Buffer, and MilliQ water. The plasmid backbone was additionally treated with FastAP Thermosensitive Alkaline Phosphatase (Sigma Aldrich) after digestion to avoid re-ligation.

For traditional restriction endonuclease cloning, the ligation reaction (10 μl) was prepared on ice by adding 1 μl of 10X Ligation Buffer, 1 μl of T4 DNA ligase, and a 7:1 ratio of insert and plasmid. The reaction was then incubated overnight at 15°C.

For the restriction free cloning protocol, the insert was amplified using primers (Key Resources Table) designed with the In-Fusion Cloning Tool (Takara-bio Inc.). Subsequently, 5 μl of the PCR product was treated with 2 μl cloning enhancer (Takara-bio Inc.). The cloning reaction was prepared by adding 2 μl of 5X In-Fusion HD Cloning enzyme mix, 1 μl linearized vector, 2 μl insert (treated with cloning enhancer), and H_2_O in a total volume of 10 μl. The reaction was incubated for 15 minutes at 50°C prior to transformation.

#### *C. elegans* plasmid generation

##### npr-11prom::WrmScarlet

*npr-11prom::WrmScarlet* was generated by cloning the *npr-11* promoter (2338bp amplified from N2 genomic DNA) into *pPD49.26. The WrmScarlet* encoding sequence *(699bp)* was amplified from pSEM90 and cloned with *NheI-KpnI*.

##### npr-11prom::nfya-1 rescue plasmid

*npr-11prom::nfya-1* was generated using the previously cloned *npr-11* promoter (2338bp amplified from N2 genomic DNA) into *pPD49.26*. The *nfya-1 cDNA* (1449bp) was amplified from N2 cDNA library and cloned into *pPD49.26* using *HindIII-BamHI*.

##### unc-47prom::nfya-1 rescue plasmid

*unc-47prom::nfya-1* was generated by cloning the *unc-47* promoter (1181 bp) from pTB169 into *pPD49.26* containing the previously cloned 1449 bp *nfya-1* cDNA using *HindIII-BamHI*.

##### unc-3prom::nfya-1 rescue plasmid

*unc-3prom::nfya-1* was generated by cloning (restriction free) the *unc-3* promoter (558bp amplified from N2 genomic DNA) into the *pPD49.26* containing the previously cloned 1449 bp *nfya-1* cDNA using *HindIII-SmaI*.

##### rab-3prom1::NLS::GFP

*rab-3prom1::NLS::GFP* was generated by cloning *rab-3prom1* (4382bp amplified from N2 genomic DNA) into the *pPD95.67* plasmid using *HindIII*.

##### rab-3prom1::nfya-1 rescue plasmid

*rab-3prom1::nfya-1 cDNA* plasmid was generated by cloning the *rab-3prom1* (4382bp amplified from N2 genomic DNA) into the *pPD49.26* containing the previously cloned 1449 bp *nfya-1* cDNA using *XbaI-SmaI*.

##### *sri-1prom::GFP* expression plasmid

*sri-1prom::gfp* was generated by cloning the *sri-1* promoter (831bp amplified from N2 genomic DNA) into *pPD95.75* using *XbaI-SmaI*.

##### *srh-277prom::GFP* expression plasmid

*srh-277prom::gfp* was generated by cloning the *srh-277* promoter (1477bp amplified from N2 genomic DNA) into *pPD95.75* using *XbaI-SmaI*.

##### *srg-32prom::gfp* expression plasmid

*srg-32prom::gfp* was generated by cloning the *srg-32* promoter (1011bp amplified from N2 genomic DNA) into *pPD95.75* using *XbaI-SmaI*.

#### Site-directed mutagenesis

The site-directed mutagenesis kit (Takara-bio Inc.) was used to alter specific cis-regulatory regions, according to the manufacturer’s protocol. First, a reporter plasmid containing the promoter and the fluorescence reporter was generated (described above). Next, the In-Fusion Cloning Tool (Takara-bio Inc.) was used to design primers to induce the desired mutation (substitution, deletion or insertion) (see Key Resources Table). After amplification, 2 μl of cloning enhancer (Takara-bio Inc.) was added to 5 μl of PCR reaction volume to remove the original plasmid. Samples were incubated at 37°C for 15 minutes, at 80°C for 15 minutes, and stored at −20°C. A 10 μl In-Fusion Cloning reaction (Takara-bio Inc.) was prepared by adding 2 μl of 5X In-Fusion HD Enzyme Premix, 1-2 μl of Purified PCR fragment and water, followed by an incubation for 15 minutes at 50°C. The samples were then transformed, plasmid DNA isolated and mutation confirmed by Sanger sequencing.

#### CRISPR-Cas9 genome editing

##### *nfyc-1(rp152)* STOP-IN cassette

A *nfyc-1* STOP-IN cassette was introduced as previously described (Wang et al., 2018). *nfyb-1(cu13)* adult hermaphrodites were microinjected with Cas9 protein, *nfyc-1* sgRNA (TTCTGGGAATTGCGACATCA), a repair template that includes a STOP-IN cassette (cttacATTATTGTCTAAAACTTCTGGGAATTGCGACGCTAGCTTATCACTTAGTCACCTCTGCTCTGGACAAACTTCCCATcacggcagatgcaagaaaaattaactttaaac), tracrRNA and *myo-2prom::mCherry.* CRISPR-Cas9 introduction of the STOP-IN cassette was detected by PCR and confirmed by Sanger sequencing. The mutant strain was backcrossed to wild-type males three times prior to analysis.

##### rab-3(rp180; syb3072) mutagenesis

Mutation of the putative NFYA-1 binding site (CCAAT>CTGCT) in the *rab-3* promoter was introduced in *syb3072(rab-3::T2A::3xNLS-gfp)* animals. Adult hermaphrodites were microinjected with Cas9 protein, *rab-3* sgRNA (AAAATATTTATGTAGGGAATTGG), a mutagenesis repair template that includes the desired mutation (GTACTCTATAAAAGATGAAACTGTAATGACTGCTTCCCTACATAAATATTTTTAACTAATTGGTCGGATA), tracrRNA and *myo-2prom::mCherry*. CRISPR-Cas9 introduction of the CTGCT mutation was detected by PCR and confirmed by Sanger sequencing. The mutant strain *rab-3(rp180; syb3072)* was backcrossed to wild-type males three times prior to analysis.

#### Behavioral assays

##### Thrash assays

To assay motility, 1-day old adult hermaphrodites were placed in 10μl of M9 liquid, allowed to recover for 10 sec. and then number of body bends was counted for one minute. A total of 10 worms were counted per each of three replicates. Animals not moving at all were censored from the experiments.

##### Locomotion assays

Locomotion speed was measured using the WormLab Imaging System (MBF Bioscience). NGM plates were seeded with 50ml OP50 bacteria and dried for 1 h. Six 1-day old adults, 20 h after mid-L4 larval stage, were picked to each assay plate and allowed to acclimate for 10 min. Videos were recorded for 1 min (3.75 frames/s). Speed was equal to distance travelled/time.

#### Calcium imaging

Animals carrying the transgene *wzIs115(rab-3prom::cameleon YC3.60)* were used to monitor neuronal activity in VNC motor neurons. Calcium imaging was performed by ratiometric fluorescence resonance energy transfer (FRET) analysis as described (Gopal et al., 2015). The fluorescent intensities of donor, acceptor and FRET channels were measured using FIJI according to the formula, corrected total fluorescence (CTF) = integrated density of cell body – (area of the cell body × mean fluorescence of background) (Gopal et al., 2021). The ratiometric FRET is calculated as the ratio between the uncorrected FRET signal intensity and the total donor intensity. Heat maps were generated using FIJI according to FRET channel intensity.

#### Fluorescence microscopy

Animals were analyzed by mounting them on a glass slide with a 5% agarose pad, using 20 mM NaN_3_ as an anaesthetic. Images were taken using an upright fluorescence microscope (Zeiss, AXIO Imager M2) and ZEN software (version 2.0).

#### Measurement of fluorescent reporter intensity

L4 animals were selected and mounted for microscopy. Differential Interference Contract (DIC) and fluorescent images of the ventral side (for PVQs) and lateral (for motor and head neurons) of the animals were captured using a 40x or 60x objective. Fluorescence was quantified using ImageJ (National Institute of Health, USA, version 1.52h). After data collection, CTCF calculation was performed with the following formula (CTCF = Integrated density – (selected area x mean fluorescence of background readings).

#### ChIP-sequencing data analysis

Publicly available NFYA-1 ChIP-seq datasets (ENCSR859SHA, ENCSR653IDD and ENCSR581THE) were obtained from modENCODE (https://www.encodeproject.org). The data was processed as per the steps outlined in the modENCODE repository for each sample. Briefly, data was processed using the recommended Illumina Data Analysis pipeline to generate aligned sequence reads. Data was then merged from each biological replicate, separated by condition (control vs experiment). These two files became the input for the PeakSeq base calling algorithm. The PeakSeq method then treats each aligned sequence read as a 200 nt fragment. The number of reads at each genomic site was counted and compared to both a randomized model of the worm genome, and the number of parallel reads obtained from sequencing the input (non-ChIP) DNA. These calculations result in an enrichment ratio and a corresponding P-value with peaks called against WS220(ce10). The average profile of peaks binding to the TSS region was generated with ChIPseeker (Yu et al., 2015). To study the distribution of peaks genome-wide, the peaks were annotated using the annotatePeak function from Homer (Heinz et al., 2010) with each peak assigned to a gene with the nearest TSS. For de novo motif discovery, default settings for motif lengths of 12 for each condition were used on the peak bed files previously generated. The identified peaks were extracted and supplied to findMotifsGenome.pl provided by Homer, using peaks as the discovery set, and the whole genome as the background. Genes that contained the NFYA motif (CCAAT) were identified using the scanMotifGenomeWide.pl command and gene enrichment tests were then performed using DAVID (Sherman et al., 2022).

## QUANTIFICATION AND STATISTICAL ANALYSIS

Statistical analyses were performed using the GraphPad Prism 9 software (graphpad.com). Statistical tests used were the t-student test (for comparing two datasets) or one-way ANOVA for multiple comparisons, and the values were expressed as mean ± standard error of the mean (SEM). Differences with a p value<0.05 were considered significant.

## AUTHOR CONTRIBUTIONS

Conceptualization, P.M., J.D. and R.P.; Methodology, P.M., P.P, J.D., A.H., S.G., D.P. and R.P.; Investigation, P.M., P.P, J.D., A.H., S.G., D.P. and R.P.; Writing – Original Draft, R.P.; Writing – Review & Editing, P.M., P.P, J.D., A.H., S.G., D.P.; Funding Acquisition, R.P.; Resources, R.P and D.P.; Supervision, R.P.

## ACKNOWLEDGEMENTS

We thank members of the Pocock Laboratory for comments on the manuscript. Some strains were provided by the *Caenorhabditis* Genetics Center (University of Minnesota), which is funded by NIH Office of Research Infrastructure Programs (P40 OD010440), and by the National BioResource Project of Japan. We thank Cori Bargmann for the *kyIs321* strain, Niels Ringstad for the *wzIs115* strain, Oliver Hobert for the following strains (*otEx6666, otEx6403, otEx233, otIs11, syb3072* and *ot1178*) and DNA (*snb-1(fosmid)::SL2::NLS::YFP::H2B; unc-11(fosmid)::SL2::NLS::YFP::H2B*), Paschalis Kratsios for *ot839* and Brent Neumann for the *WrmScarlet* plasmid. This work was supported by the following grants: NHMRC (Senior Research Fellowship GNT1137645 to R.P.) and veski Innovation Fellowship (VIF23 to R.P.).

