## Supplementary figures and images for "Nuclear Factor-Y is a Pervasive Regulator of Neuronal Gene Expression"

### Figure S1

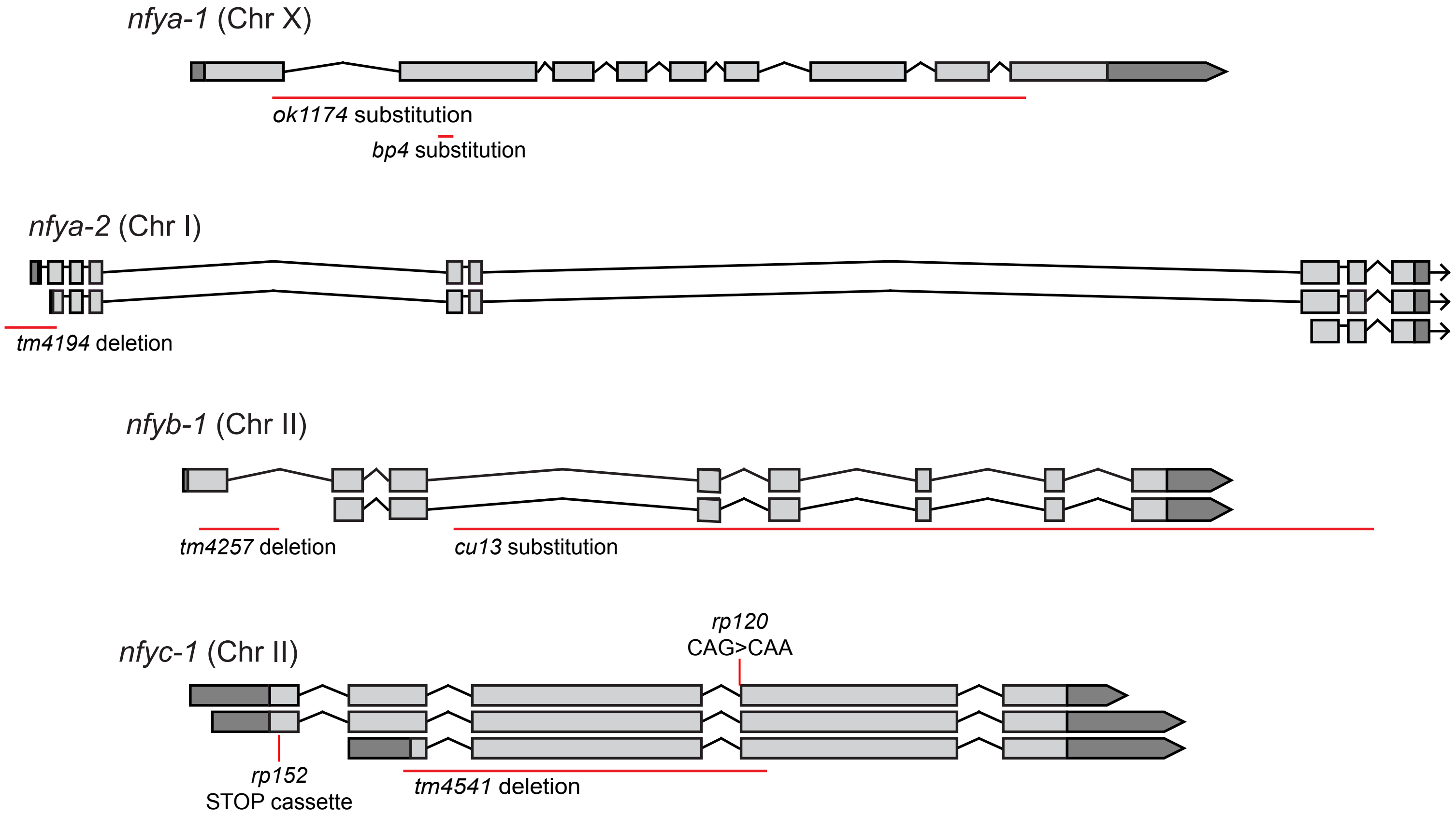

### Figure S2

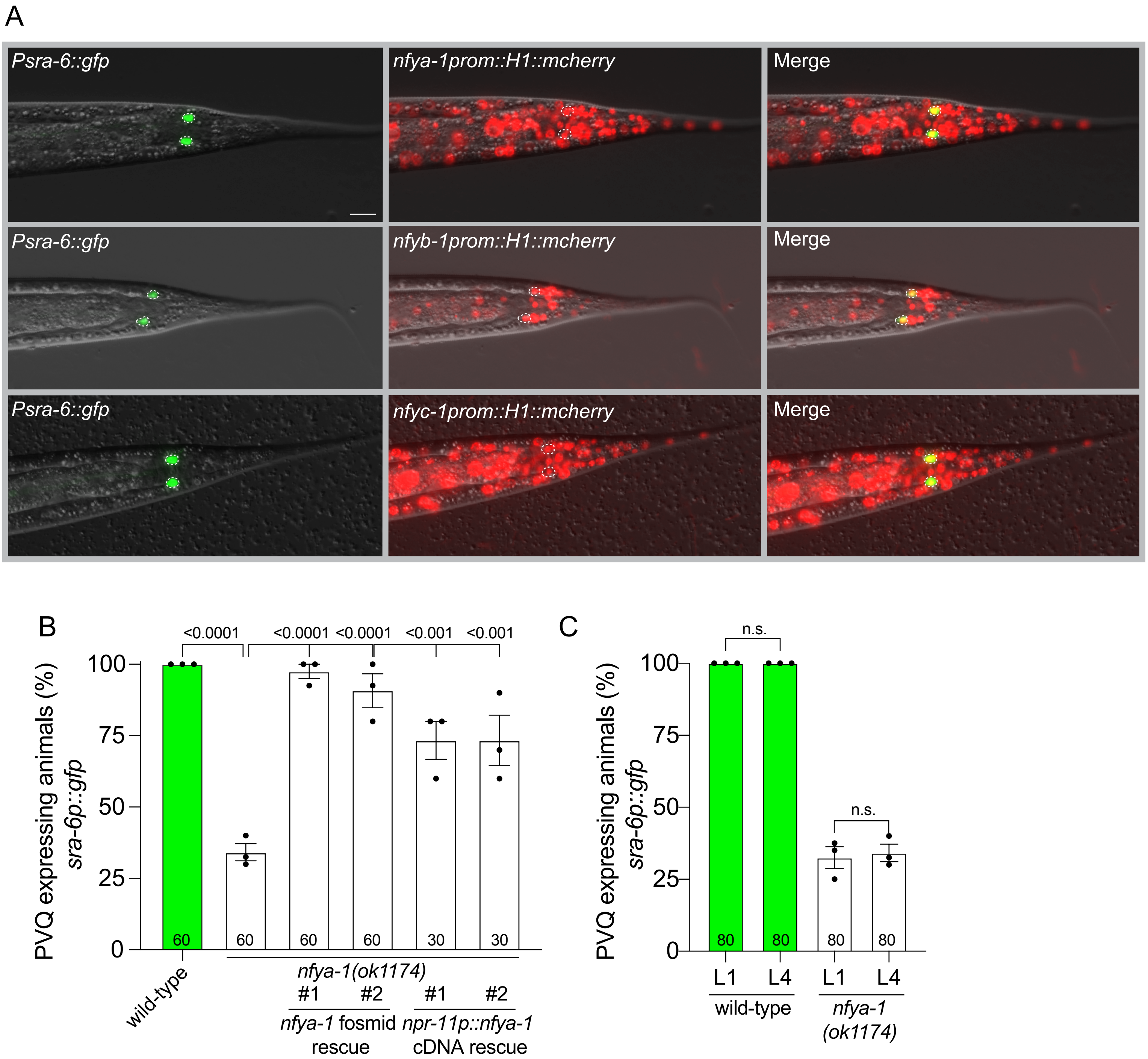

### Figure S3

A

| Gene | CCAAT sites | ATTGG sites | Promoter length in reporter (bp) |
|---|---|---|---|
| *sra-6* | 7 | 4 | 3775 |
| *srg-32* | 0 | 1 | 1041 |
| *srh-277* | 3 | 0 | 1477 |
| *sri-1* | 1 | 2 | 831 |
| *dop-1* | 7 | 5 | 2009 |
| *egl-47* | 3 | 7 | 4851 |
| *npr-11* | 1 | 4 | 2388 |

B

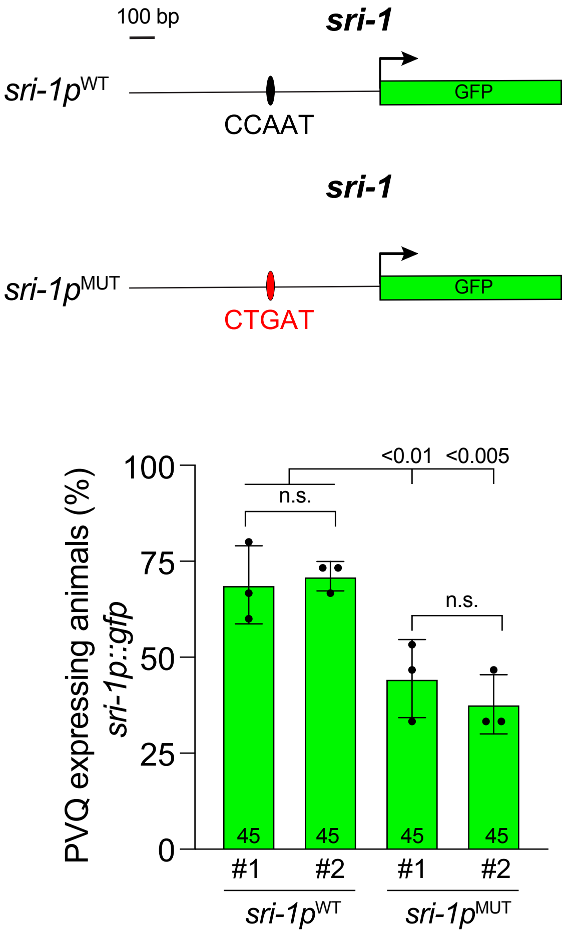


C

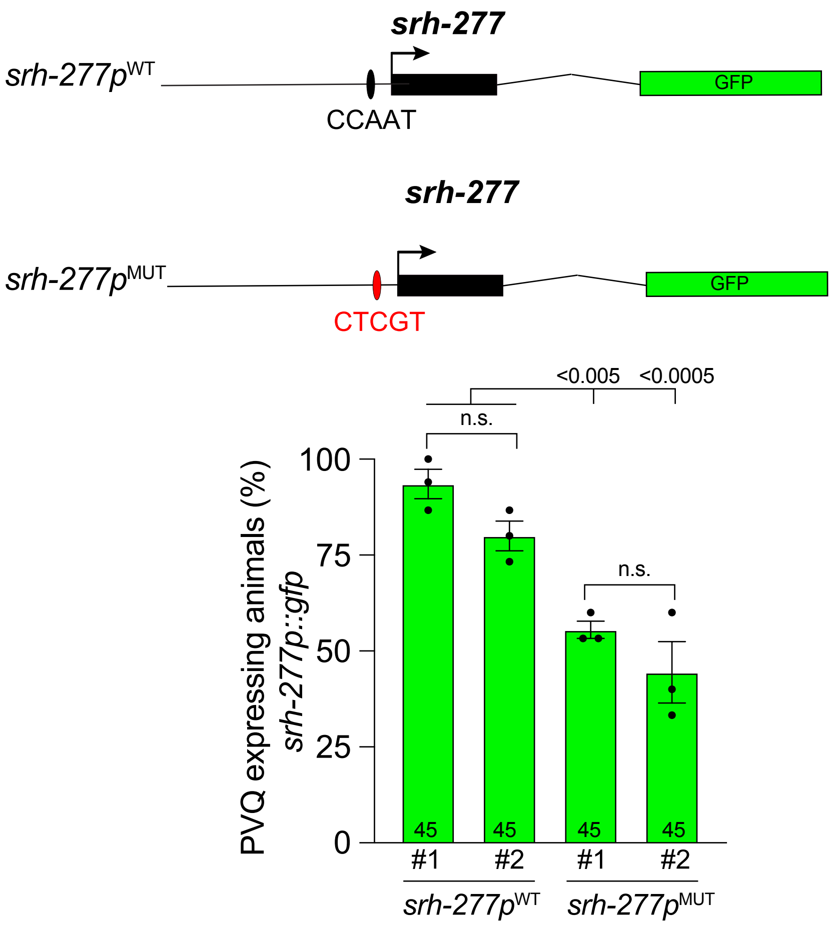

### Figure S4

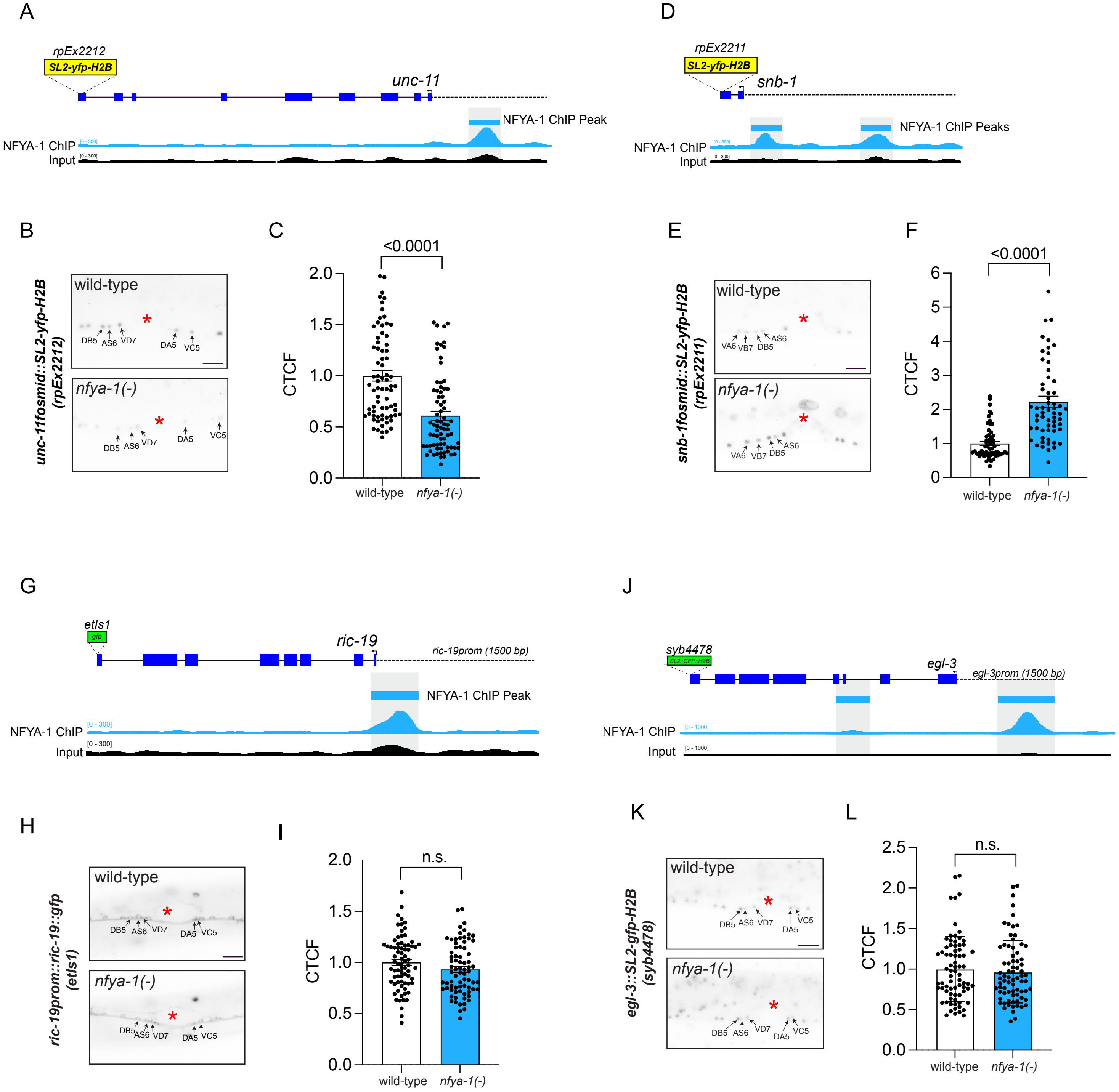

### Figure S5

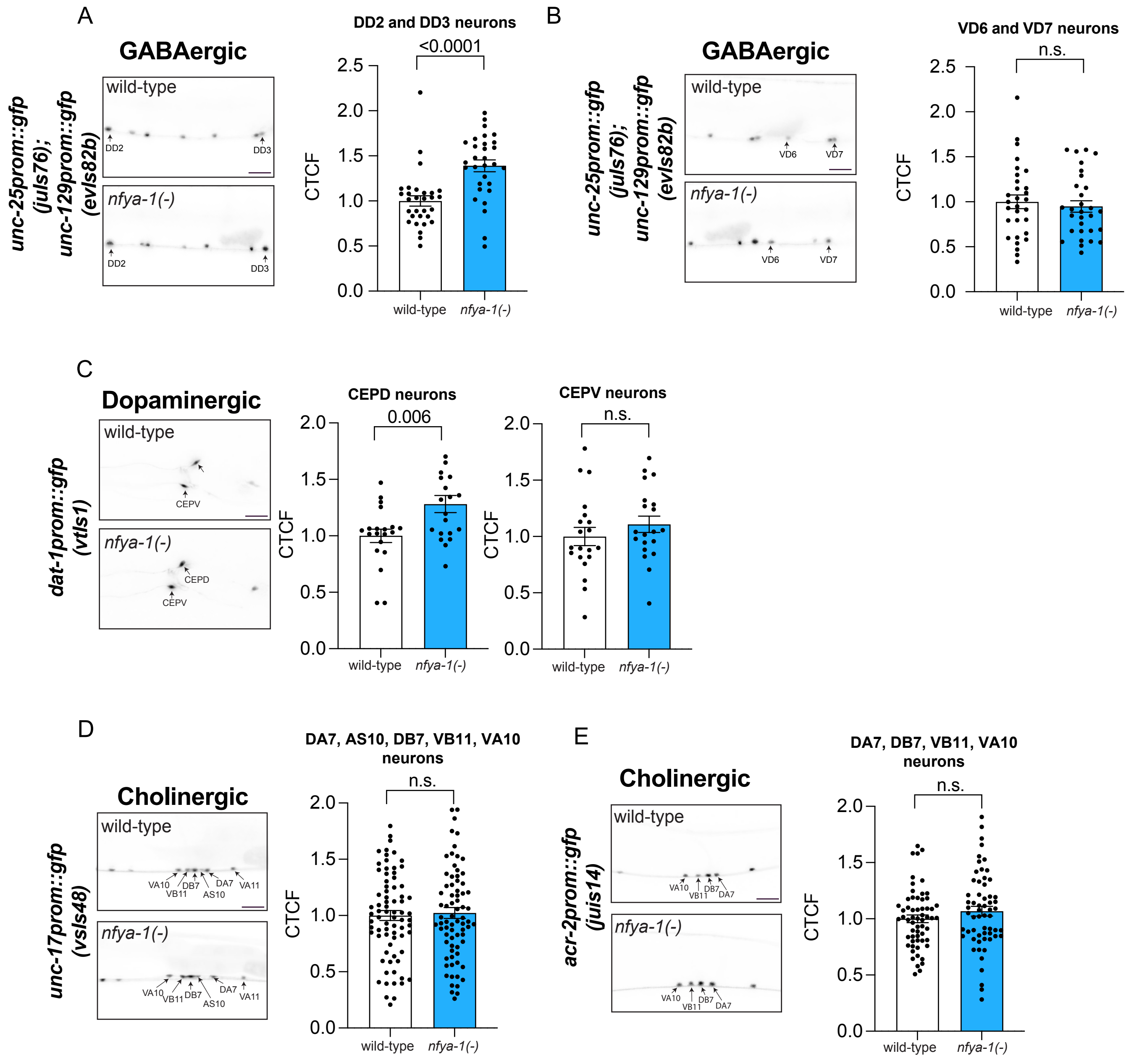
